# Evidence for the plant recruitment of beneficial microbes to suppress soil-borne pathogen

**DOI:** 10.1101/2020.07.31.231886

**Authors:** Hongwei Liu, Jiayu Li, Lilia C. Cavalhais, Cassandra Percy, Jay Prakash Verma, Peer M. Schenk, Brajesh Singh

**Author notes:** These authors contributed equally (Peer M. Schenk, Brajesh Singh).

## Abstract

- Emerging experimental framework suggests that plants under biotic stress may actively seek help from soil microbes, but empirical evidence underlying such a ‘cry for help’ strategy is limited.
- We used integrated microbial community profiling, pathogen and plant transcriptive gene quantification and culture-based methods to systematically investigate a three-way interaction between the wheat plant, wheat-associated microbiomes and *Fusarium pseudograminearum* (*Fp*).
- A clear enrichment of a dominant bacterium, *Stenotrophomonas rhizophila* (SR80), was observed in both the rhizosphere and root endosphere of *Fp-*infected wheat. SR80 reached 3.7×10^7^ cells g^-1^ in the rhizosphere and accounted for up to 11.4% of the microbes in the root endosphere. Its abundance had a positive linear correlation with the pathogen load at base stems and expression of multiple defense genes in top leaves. Upon re-introduction in soils, SR80 enhanced plant growth, both the below- and above-ground, and induced strong disease resistance by priming plant defense in the aboveground plant parts, but only when the pathogen was present
- Together, the bacterium SR80 seems to have acted as an early warning system for plant defense. This work provides novel evidence for the potential protection of plants against pathogens by an enriched beneficial microbe via modulation of the plant immune system.

## Introduction

Plants have been subject to selective pressures from pathogens, pests, and undesirable soil (e.g., nutrient deficient) and weather (e.g., drought) conditions since their evolutionary origin (Chakraborty & Newton, 2011; Liu *et al.*, 2019). Assemblages of host-specific microbiomes in the rhizosphere, root endosphere and other niches, such as the phyllosphere (leaf) and anthosphere (flower) have been reported (Edwards *et al.*, 2015; Álvarez-Pérez *et al.*, 2019; Grady *et al.*, 2019). Emerging evidence indicates that these microbial symbionts, rather than merely acting as tenants, intimately interact with plants and influence their immune systems and multiple processes of plant growth and development (Liu *et al.*, 2019). However, detailed mechanistic evidence for these interactions is missing and how these hugely diverse commensal microbes interact with plants is largely unknown. Addressing these knowledge gaps is urged to support future translational research for the application of microbe-based products in sustainable agriculture.

The rhizosphere is a hotspot for plant-soil-microbe and microbe-microbe interactions. The microbes and their interactions can largely prevent pathogen outgrowth and extend the plant capacity for disease resistance (Durán *et al.*, 2018). Recent studies revealed that some disease-resistant crop varieties (e.g. tomato and common bean) enrich particular sets of bacterial species in the rhizosphere to suppress pathogens (Kwak *et al.*, 2018; Mendes *et al.*, 2018). This suggests that specific soil microbes/microbial functions contribute to protection against plant diseases. Furthermore, it has been shown that diseased plants significantly promoted specific beneficial microbes in their associated microbiomes, which could act as an extended layer of plant defense and a legacy to enhance survival rates of their offspring (Berendsen *et al.*, 2018). However, whether the rhizosphere and root endophytic microbiota are coordinated to modulate plant-pathogen interactions and plant performance is unknown.

Recent theoretical framework of co-evolution also suggests that recruitment of beneficial microbes upon biotic stress is likely a survival strategy conserved across the plant kingdom although empirical evidence is still scarce and mechanistic pathways remain poorly defined (Liu & Brettell, 2019; Liu *et al.*, 2019). Whether such a cry for help strategy applies to crop-pathogen interactions in field conditions is unknown. Moreover, plants differ in physiology and immune response to pathogen invasions, meaning mechanisms are likely to be plant species-dependent. In this study, we investigated interactions between durum wheat (*Triticum turgidum var. durum*) and its microbial symbionts in the rhizosphere and root endosphere upon infection with the fungal pathogen *Fusarium pseudograminearum* (*Fp*). Crown rot (CR) disease caused by *Fp* is a devastating soil-borne disease, which can infect winter cereal crops of all stages and lead to large losses in global cereal production (estimated at $79 million/year in Australia alone) (Akinsanmi *et al.*, 2004). However, so far, there are no CR-resistant crop varieties and effective control through chemical fungicides is currently not available. Using wheat-*Fp* system, we aimed to explore responses of plant microbiomes to pathogen infection and identify mechanisms underpinning plant-microbiome interactions that may mediate disease resistance of the plant.

In an effort to achieve our aims, durum wheat plants were sampled from a field experiment where *Fp* inoculations have been historically occurring at the Queensland Department of Agriculture and Fisheries (DAF) research farm at Wellcamp (Australia). Soil and tissues from symptomatic plants that were naturally infected with *Fp* were collected, along with those from asymptomatic plants, to identify key microbial taxa in the plant microbiome that are associated with plant defense responses. We profiled microbial communities in the rhizosphere and roots using bacterial and fungal amplicon sequencing. Plant defense status was evaluated by determining the transcript abundances of defense genes involved in the salicylic acid (SA) and jasmonic acid (JA) signaling pathways. Bacteria were then isolated from *Fp*-infected plants and effects of a disease-enriched bacterium on durum wheat defense and growth were investigated in glasshouse experiments.

## Methods and Materials

### Fp treatments in field experiments and sampling

Field samples were collected from a wheat trial conducted in 2015 at the Queensland Department of Agriculture and Fisheries (DAF) Experimental Farm at Wellcamp (27°33’54.7”S, 151°51’52.0”E), Queensland, Australia. The soil is a black Vertosol with chemical properties listed in Table S1. Farm management histories including past yield trials, CR trials and fallow periods are shown in Table S2. Plus and minus *Fp* (the causal agent for CR disease) inoculated yield plots (6 m x 2 m) were sown in a randomized complete block design in June 2015. Colonized millet grain inoculum (Percy *et al.*, 2012) was applied in the furrow in a band above the seed at planting. Experimental plots were surrounded by a 7-row buffer zone (2 m each) of non-inoculated durum wheat (Jandaroi variety), running 150 m along the length of the experiment. CR was observed in many plants in the non-inoculated Jandaroi buffer during tillering and as jointing progressed. Diseased plants were scattered amongst asymptomatic plants. *Fp*-infected plant residues remaining from a previous CR experiment (2010) may have been retained in the field and continued to provide a source of inoculum to infect subsequent hosts. The research presented in this study compares samples taken from plants that were asymptomatic and symptomatic for CR in Jandaroi buffer rows, as we aimed to investigate plants naturally infected with *Fp*.

Thirteen weeks after sowing, both healthy and infected plants were carefully uprooted using a shovel and separated into independent individuals from three different locations of the field in September 2015. Forty healthy and 18 infected plants were collected for microbiome analyses. The top ∼10 cm of two to three leaves were cut, transferred into a 15 mL Falcon tube and immediately frozen in dry ice. The leaf along with stem and root (soil attached) samples were stored in dry ice and transported to the laboratory on the same day and preserved at -80°C. Approximately 5 grams of fresh roots (rhizosphere soil attached) collected from a *Fp*-infected plant were kept at 4°C until microbial isolation.

### Processing of the rhizosphere soil and root samples

Stems, roots and soils were separated before further processing. The procedures included three steps. (i) The most basal internode/base stem (∼5 cm that covers an area of brown discoloration of each plant) was cut and scored for disease levels following the protocol below. The stem samples were ground into a fine powder in liquid nitrogen and stored at - 80°C for DNA extraction. (ii) Bulk soil was removed from roots by shaking them vigorously. (iii) Rhizosphere soil was then separated from roots by shaking root and soil in 25 mL 0.1□M sterile phosphate buffer (7.1□g Na_2_HPO_4_, 4.4□g NaH_2_PO_4_·H_2_O added to 820□mL deionized water, pH 7.0) in a 50 mL plastic conical centrifuge tube at 200 rpm for 5 min. Roots were then transferred to a new tube for subsequent procedures. The soil suspension was centrifuged at 4,000 g for 15 min and the obtained soil pellet was regarded as the rhizosphere soil and stored at -80°C until genomic DNA extraction. Roots were thoroughly washed under distilled water, followed by surface sterilization to remove microbes on the surface by shaking roots in 25 mL of 4% sodium hypochlorite solution at 200 rpm for 5 min. The roots were then washed three times with sterile phosphate buffer. The last wash was inoculated on a nutrient agar plate incubated for 2 days at 37°C and no viable colonies formed, which indicated that the disinfection procedure was efficient. Roots were air-dried in a laminar hood and stored at -80°C until grinding step in liquid nitrogen leading to a fine powder for DNA extraction.

### Disease scoring

CR was rated based on the percentage of brown discoloration at the stem base using a 0 to 4 scale previously reported (Wildermuth & McNamara, 1994). A score of 0 indicates no visible discoloration, and a score of 1, 2, 3, and 4 describe discoloration of 0∼25%, 26% to 50%, 51%∼75% and 76%∼100%, respectively.

### DNA extraction from soil and plant samples

Genomic DNA (gDNA) was extracted from about 0.2 g plant (stem or root) samples using a Maxwell^®^ 16 LEV Plant DNA Kit on a Maxwell^®^ 16 Instrument (AS2000) according to the manufacturer’s instructions. Soil gDNA was extracted from 0.25 g soil per sample using the PowerSoil™ DNA Isolation kit (MO BIO Laboratories, Carlsbad, CA) as per the manufacturer’s recommendations. Plant and soil DNA concentrations were determined using a Qubit™ fluorometer with Quant-iT dsDNA HS Assay Kit (Invitrogen). The DNA samples were stored at -20°C until further analysis.

### Fp and total fungi quantification

*Fp* and total fungi colonizing the plant and soil samples were quantified using qPCR. The details of the qPCR system and thermal conditions are described in the Supplementary Materials of this study. The proportion of *Fp*/total fungi DNA to wheat DNA was counted as the relative abundance of *Fp*/fungi in wheat plants (Melloy *et al.*, 2010). *Fp* and fungi were estimated using (i) the *Tri5* gene from the trichothecene cluster responsible for trichothecene production of *Fusarium* species, and (ii) the fungal ribosomal 18S rRNA gene, respectively (Melloy *et al.*, 2010). Wheat actin-binding protein coding sequence was used as the reference gene (Table S3). Quantification of these genes was performed using SYBR Green on a ViiA™ 7 sequence detection system (Applied Biosystems, USA) with two replications for each sample. Amplification specificity for each gene was examined by running melt curve analysis, and the *Fp* amounts were then estimated according to

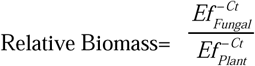

where *Ef* is PCR amplification efficiency, which was calculated by LinRegPCR 7.5 based on raw PCR amplification data (Ramakers *et al.*, 2003). The relative abundance of *Fp* in soil was also calculated, by comparing Ct values of each sample against a standard curve that was generated by qPCRs using serially diluted *Tri5* amplicons.

### Quantification of defense-related genes in the leaf

RNA was isolated from ground leaf samples using Maxwell^®^ 16 Total RNA Purification (Promega) kit on an automated Maxwell^®^ 16 Instrument as per the manufacturer’s recommendations. RNA concentrations were quantified using a Qubit™ fluorometer (Invitrogen). cDNA was then synthesized from 2.5 μg RNA in a 20 μL reaction using both random hexamers and oligo(dT) primers provided with the Tetro cDNA Synthesis kit (Bioline™). Ten defense genes were quantified using the SYBR Green qRT-PCR kit on a ViiA™ 7 sequence detection system. These genes are marker genes for JA and SA signaling pathways, characterized in our previous study (Table S3) (Liu *et al.*, 2016). The 18S rRNA gene was used as the housekeeping gene for normalization, and the cDNA was diluted to 1: 500 before the amplification of this gene. The reverse transcriptase qPCR (qRT-PCR) system and thermal conditions are described in the Supplementary Materials.

### Plant and soil microbial community profiling using amplicon sequencing

The universal primers 926F (Engelbrektson *et al.*, 2010) and 1392R (Peiffer *et al.*, 2013) were used for the amplification of bacterial and archaeal 16S rRNA genes in soil samples (Table S3). The primers 799F and 1193R that preferentially amplify archaeal and bacterial DNA in plant materials were used to amplify root endophytic microbes (Table S3). B adaptor was linked to a key (TCAG) and connected to the above template-specific forward primers. A adaptor was linked to a key sequence (TCAG) and a sample-specific molecular identifier, and was then connected to a template-specific reverse primer (Table S3). Amplifications were performed in duplicates for each sample and combined, using the PCR system and conditions described in the Supplementary Materials. PCR products were examined using agarose gel electrophoresis, and microbial amplification products for root DNA samples were recovered from gels to remove the plant mitochondrial DNA-derived amplicons which co-amplify using the 799F and 1193R primers but are of a different length. The obtained 16S rRNA amplicons (∼400□bp) were purified using Agencourt AMPure XP beads (Beckman Coulter, Inc.) and quantified with a PicoGreen dsDNA Quantification Kit (Invitrogen). Amplicons were then dual indexed using the Nextera XT Index Kit (Illumina) according to the manufacturer’s instructions. Equal concentrations of each indexed sample were pooled and sequenced on an Illumina MiSeq as per the manufacturer’s instructions at the University of Queensland’s Institute for Molecular Biosciences (UQ, IMB). Additionally, the fungal amplicon sequencing for soil samples was performed at the Western Sydney University using the primer sets FITS7 (Ihrmark *et al.*, 2012) and ITS4 (Glass & Donaldson, 1995) as per the standard MiSeq sequencing procedures. Bioinformatic analyses are described in the Supplementary Materials of this study.

### Bacterial isolation

One gram of roots combined with rhizosphere soil from *Fp*-infected wheat were cut into small pieces and homogenized with autoclaved glass beads (710-1180 µm, Sigma) in a 2 mL centrifuge tube for 3 min using a TissueLyzer (Qiagen). The obtained mixture was serially diluted in 5.0 mL sterile phosphate buffer and 150 µL aliquots of 10^−4^ to 10^−7^ dilutions were spread onto plates containing a range of media: nutrient agar (NA), Pikovskaya’s agar, tryptophan soy agar, King’s B agar and *Stenotrophomonas* spp. selective medium (32 mg L^-1^ Imipenem added, Adooq Biosciences) (Bollet *et al.*, 1995). Plates were incubated at 30°C in an incubator (Thermoline Scientific) for 2-5 days. Bacterial colonies were then picked based on morphology, color and margin, and further purified by streaking on new NA plates. Purified strains were stored in 25% glycerol at -80°C and sub-cultured on NA plates for the required analyses.

### Antifungal assays

*Fp* strains CS3427, CS3321, CS3096 (Chakraborty *et al.*, 2010) and A11 (strain in the field trial) were used for this assay. They were cultured on potato dextrose agar (PDA) to a whole plate size, and a mycelial plug of each fungus was placed in the center of a dual media plate (½ PDA and ½ NA media). Tested bacteria were streaked at four ends of the plate and cultured at 28°C for 7 days to examine antagonistic effects on these *Fp* strains. Two *Burkholderia* bacterial strains were used as positive controls, and three replicates were performed for the assay. Plates that inoculated with *Fp* only served as negative controls.

### Bacterial identification and whole genome sequencing of SR80

Bacterial strains were identified based on their 16S rRNA gene sequence. They were cultured in nutrient broth for 24 h, pelleted by centrifugation, washed and re-pelleted in PBS buffer twice. Bacteria were then subjected to gDNA extraction using the DNeasy PowerSoil Pro kit (Qiagen). A region of the 16S rRNA gene was amplified using 27F and 1492R primers (Table S3) and sequenced on a capillary sequencer at the Hawkesbury Institute for the Environment (HIE). Sequences were quality-checked, trimmed and BLAST-searched against the nucleotide database of NCBI using Geneious 10.1.3. Sequences of *Stenotrophomonas* isolates were deposited in the NCBI database under GenBank accession numbers MT151295 to MT151301. The identification of different *Stenotrophomonas* spp. was further examined by BOX-PCR (for conditions see Supplementary Materials). Sequences of other bacterial isolates are available under MT158490-MTI158578. Whole genome sequencing of SR80 was conducted using the Illumina MiSeq platform by Guangzhou Magigene Technology Co., Ltd. Sequencing success and quality were checked with FastQC (0.11.8). A total of 11,331,006 high quality reads (Q>35) were obtained. De novo assembling for high quality scaffolds was performed using SPAdes v3.9.0 (Nurk *et al.*, 2017). The assembled sequences were deposited in the NCBI database under GenBank accession number SAMN14299480. Hereafter, the components of coding genes, non-coding RNA (ncRNA), and functional annotation using a range of databases including NR, Swissprot, COG, KEGG, GO were performed. A circular map of the genome was obtained using Circos version 0.69 (Krzywinski et al., 2009). The prediction of non-encoding 23S rRNA and 16S rRNA were performed using rRNAmmer 1.2 (Lagesen *et al.*, 2007).

### Glasshouse experiments to assess effects of SR80 on wheat growth and defense

The biological relevance of SR80 on wheat was investigated in glasshouse experiments. The soil used for wheat cultivation was collected from the Wellcamp site in August 2019 and autoclaved twice to ensure *Fp* elimination. Three treatments including *Fp* only, SR80 only, *Fp* & SR80, and control were applied in pot experiments at sowing time (8×12 cm pots, approx. 200 g soil each). The *Fp* inoculum grown on whole wheat grain was developed using the strain *Fp* A11#04, which was ground, sieved (2 mm) and stored in low humidity at 4°C until use. *Fp* was applied using an inoculation method modified from Wildermuth and McNamara, 1994 (Fig.S1). Briefly, six Jandaroi seeds were sown in soil and covered with a thin layer of soil, and topped by a layer of the *Fp* inoculant (1.0 g) in the form of fine powder.

For treatments that excluded *Fp*, the autoclaved *Fp* inoculant was applied to reproduce the chemical environment but the pathogen was not present in a living form. For bacterial treatments, 40 mL of the SR80 suspension (in 50 μM PBS buffer, pH 7.0, OD_600_=1.0) was inoculated to each pot, which provided ∼2.4×10^8^ SR80 cells g^-1^ bulk soil. As control, the same amount of PBS buffer was added to treatment groups that excluded bacterial inoculations. Soils in pots were then watered to field capacity to allow seeds to germinate. Thereafter, soil was watered daily. The height of seedlings that emerged 3 days after sowing was recorded twice daily (10 am and 4 pm) for 15 days. Disease and plant survival rates were recorded once per day. The diameter of the stem base was recorded on the 7^th^ and 14^th^ day after seedling emergence. Approx. 1.5 g of bulk soil was collected from the soil surface from each pot at the 5^th^, 10^th^ and 15^th^ day for determining the abundances of SR80 and *Fp*. When harvested, wheat shoots were cut, weighed, wrapped in aluminum foil and stored at - 80°C for total RNA extraction. Roots were carefully separated from soils, thoroughly rinsed with tap water, air-dried on paper tissues and weighed. Due to total lethality of plants in the *Fp*-treated control group, an additional experiment was conducted, which included the *Fp* and *Fp*+SR80 treatments using the same method as above but with a lower inoculant load per pot (0.6 g). This provided sufficient plant material for RNA extractions and qRT-PCR analyses.

### Statistical analyses

Statistical analyses were implemented in R3.6.1. Linear model (Pearson correlation) was performed using the package ggpubr (0.1.6) to determine (i) correlations of defense gene abundance with disease severity, and (ii) correlations of SR80 abundance in soil and roots with *Fp* amounts in plants. Data transformation was performed where needed to ensure data normalization before conducting statistical analyses. The effects of *Fp* and SR80 on responses of soil and plant microbial communities were investigated by permutational multivariate analysis of variance (PERMANOVA) using Hellinger transformed OTU abundances. This analysis was performed using the package vegan 2.5-6 (Dixon, 2003).

Treatment effects on wheat growth, defense gene expression, alpha diversity of microbial communities, and the abundance of SR80 and *Fp* were analyzed using one-way ANOVA and Tukey post hoc tests.

## Results

### Plants were naturally infected by Fp and defense signaling pathways were activated in the field study

Fifty-eight plants were collected from a field trial at the Wellcamp site. Symptomatology inspections revealed that 40 of these plants were asymptomatic for CR and 18 plants exhibited CR symptoms at different severity levels (Fig.1a). Disease severity of all plants was visually rated based on discoloration of the stem and the level of *Fp* infection was quantified by real-time quantitative polymerase chain reaction (qPCR) targeting the *Tri5* gene (Fig.1b). The two methods provided consistent results (Fig.1b), as plants with visible CR symptoms also had higher pathogen load quantified by qPCR (relative abundance ≥1) than the asymptomatic plants (Fig.1a). Moreover, *Fp* load had a linear correlation with the total fungi in base stem tissues (Fig.S2), indicating that *Fp* was the main component driving the increases of fungal abundance in CR-infected plants in the field. Genes involved in plant defense signaling pathways including *PR2* (a beta-1,3-endoglucanase), *PR3* (a chitinase), *PR4a* (wheatwin), *PR5* (a thaumatin-like protein), *PR10* (a wheat peroxidase), *TaPAL* (phenylalanine ammonia lyase) and *Lipase* were highly expressed in top leaves of *Fp*-infected plants (Fig.1c-j). These genes are known to be markers for the activation of JA and/or SA signaling pathways in wheat (Liu *et al.*, 2016). A positive linear correlation was observed between transcript abundances of these defense genes and *Fp* load in the stem tissues (Fig.1c-j, m&n). In contrast, the expression of two genes, *TaNPR1* (wheat nonexpressor of pathogenesis-related genes 1) and *WCI3* (a sulfur-rich/thionin-like protein), were not influenced by *Fp* (Fig.1k&l). Collectively, JA and SA defense signaling pathways were activated in wheat by *Fp* (Fig.1m&n), which indicates successful progression of CR and activation of plant defenses in the field.

**Fig.1.**
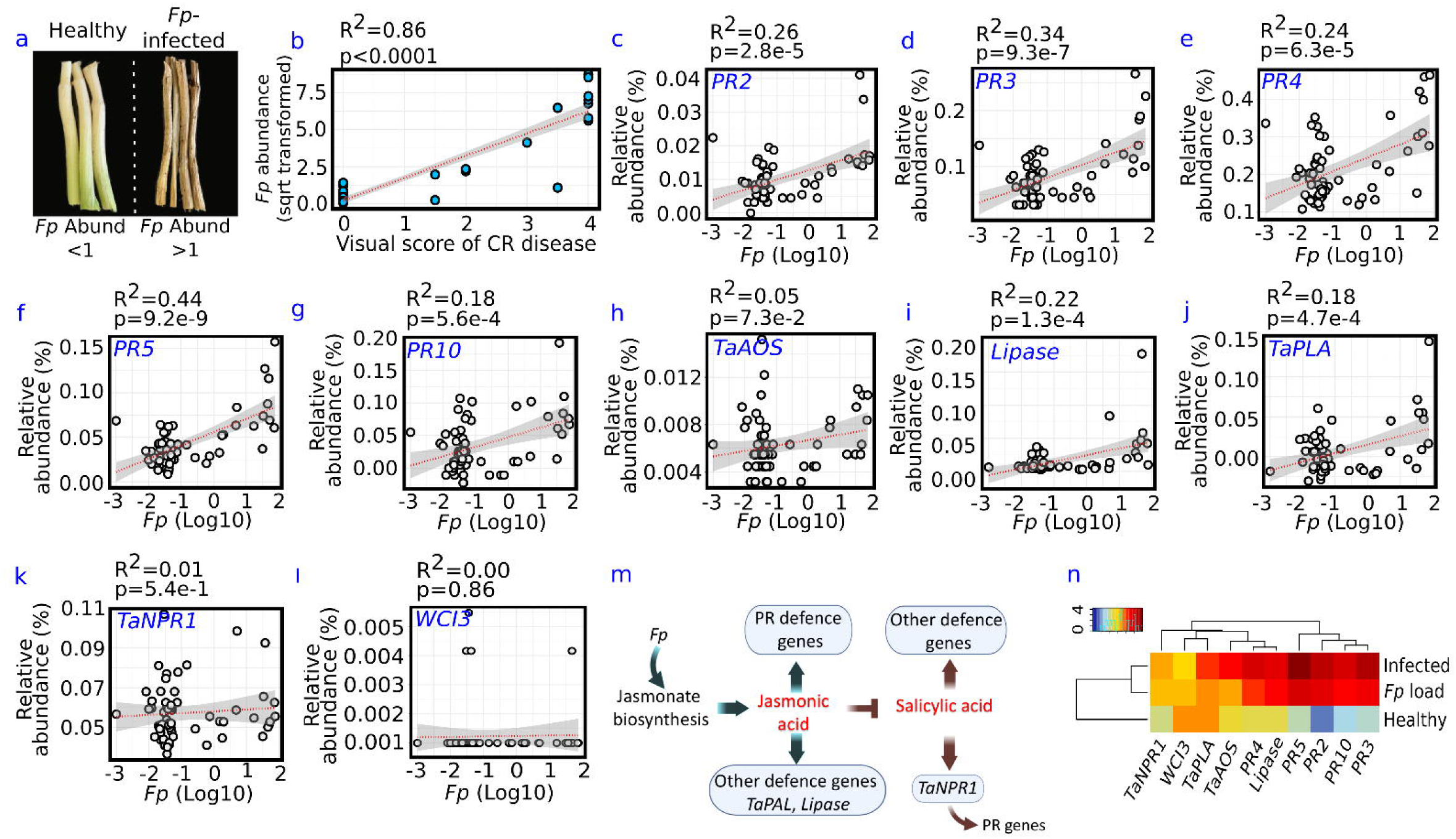
The effects of *Fp*-infection on wheat stem and expression of defense genes. An example of healthy and *Fp*-infected wheat stems (internode 1) (**a**). *Fp*-infected wheat stems had a brown discoloration (relative abundance of *Fp*≥ 1) while the healthy stems were white/light green (*Fp* abundance< 1). A Pearson correlation of *Fp* abundance with visually rated disease levels at stem base (b). Effects of *Fp* infection on the transcription of ten genes associated with jasmonic acid (JA) and salicylic acid (SA) signaling pathways in wheat (c-l). Linear correlation tests were performed for these defense genes with the *Fp* load present in the stem base. Each grey circle represents an independent measurement (*n* = 58). Red dashes show significant regression and the overlaid shaded area represents 95% confidence intervals. Both the JA and SA signaling pathways were activated by the *Fp* infection (m), and a heatmap summarizing the correlation of the expression of defense genes in leaves with the disease severity in wheat (n).

### Microbiome structure differed between healthy and diseased wheat

Wheat roots seem to have acted as a barrier to select soil microbes, resulting in phylogenetic conservation in the rhizosphere and root endosphere (Fig.S3a). These root-associated compartments selected microbial phylotypes within the Proteobacteria, Actinobacteria and to a lesser extent Bacteroidetes and Firmicutes while Acidobacteria, Gemmatimonadetes and Archaea were almost depleted from the root endosphere (Fig.S3a). Meanwhile, a gradient decrease of alpha diversty (e.g. observed species) from bulk soil to the root was observed (Fig.S3b). However, no differences were seen in the alpha diversity for the rhizosphere and root endosphere microbial communities between the healthy and diseased plants (Table S4).

The microbial composition in the rhizosphere of *Fp*-infected plants significantly differed from that of healthy plants (PERMANOVA, *P*=0.0002) and were marginally different in the root endosphere (*P*=0.073) (Fig.2a&b). Furthermore, the rhizosphere soil fungal community composition significantly differed between healthy and diseased plants (*P*=0.001) (Fig.S4). In the redundancy analyses for the rhizoshere communities (RDA, Fig.2a), bacterial OTU 89589 is strongly correlated with the *Fp*-infected plants and is one of the major dominant OTUs in the separation of the diseased plants from healthy plants in the primary axis of the RDA (RDA1), well separated from rest of the OTUs. A consistent pattern was also found in the root endosphere, where OTU 41442 was among the most discriminating OTUs for infected plants (Fig.2b). Further analyses revealed that these two OTUs had 100% nucleotide similarity within the 16S rRNA region amplified, indicating that they were the same bacterial strain (Fig.S5a, b), and belonged to *Stenotrophomonas* spp.

**Fig.2.**
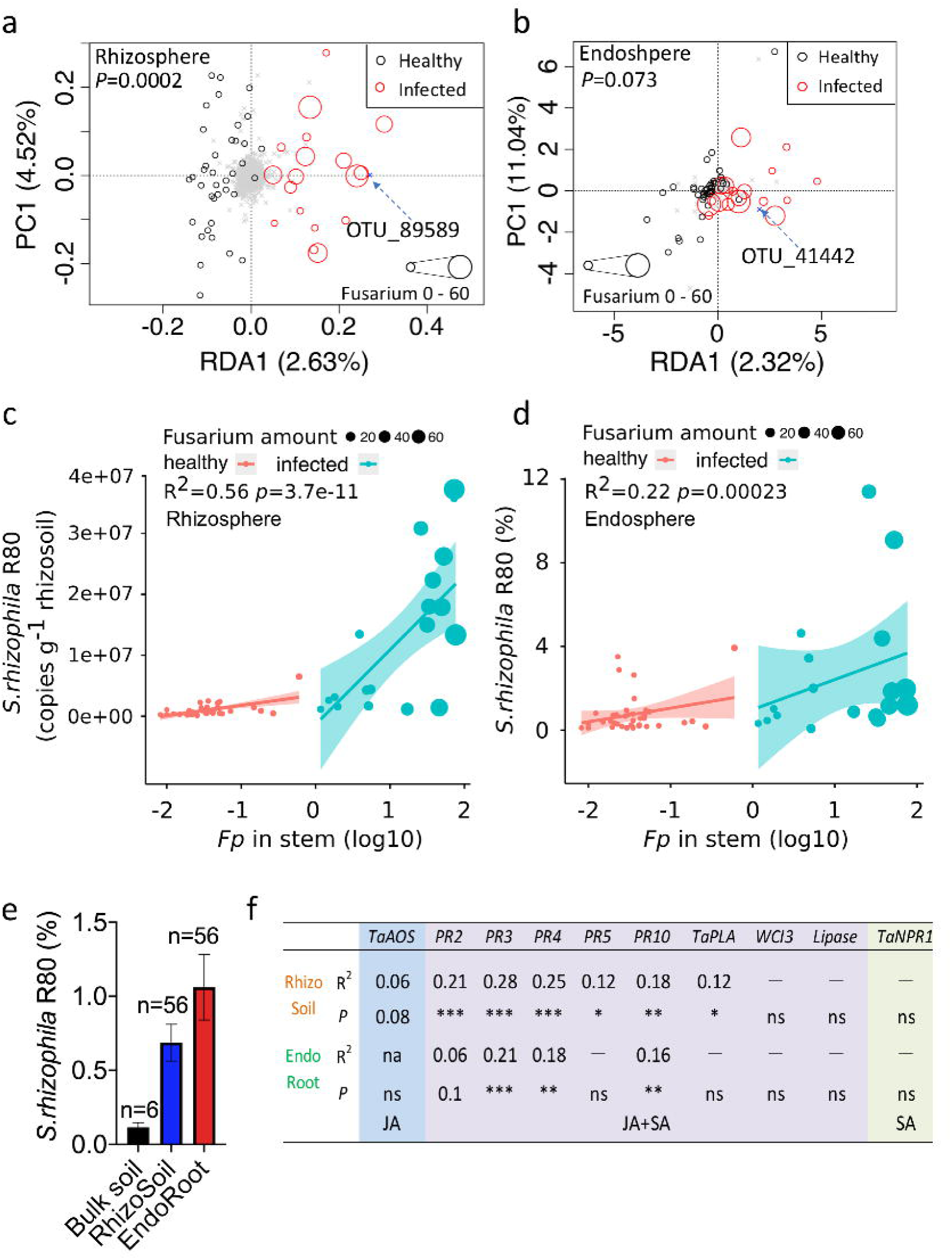
Effects of *Fp* infection on wheat-associated microbial communities. Redundancy anaysis sumarizing variations in composition of microbial communities in the rhizosphere soil (a) and root endosphere (b) attributed to crown rot. Grey crosses shown in (a) and (b) represent bacterial OTUs detected in the rhizosphere and root endosphere. Pearson correlations of the SR80 abundance in the rhizosphere soil (c) and root endosphere (d) with *Fp* abundance in wheat stems. Gradient increases in relative abundance of SR80 from the bulk soil to the rhizosphere, and to the root endosphere (e). A table summarizing the correlation of SR80 abundances with defence gene expression in leaves (f). Asterisks indicate significant correlations (* *P*□<□0.05, ** *P*□<□0.01, *** *P*□<□0.001).

Further analyses revealed that the abundance of OTU 89589 (in the rhizosphere) positively correlated to the *Fp* load in the stem tissues of infected plants (relative abundance ≥ 1) (Fig.2c). This was less noticeable in asymptomatic plants (relative abundance <1) (Fig.2c). *Fp* infection had contributed to the enrichment of this bacterium to as much as 3.7×10^7^ cells per gram of the rhizosphere soil (Fig.2c). Similarly, the abundance of this bacterium in the root endosphere also positively correlated with *Fp* abundance, accounting for up to 11.4% of the bacterial community in roots (Fig.2d). These results suggest that the wheat-associated microbiome had significantly changed in composition in response to *Fp*-infection, and specifically enriched for a *Stenotrophomonas* spp. in the process. We also observed that selection for the bacterium seems to have occurred even when wheat plants were healthy (Fig.2e). Interestingly, the abundance of this bacterium also positively correlated with transcript abundances of all PR defense genes tested (Fig.2f), suggesting that the bacterium is likely to have contributed to the activation of plant defenses in the field.

### Bacterial isolation, identification and whole genome sequencing

We attempted to isolate *Stenotrophomonas* spp., to investigate its biological implications on wheat performance via controlled *in vivo* and *in vitro* tests. A total of 179 bacterial isolates were recovered from the *Fp*-infected plant rhizosphere and roots using different media, including a selective medium for *Stenotrophomonas* spp. Among these, seven isolates were identified as *Stenotrophomonas* spp. by 16S rRNA Sanger sequencing, and these belonged to four different strains (identified using BOX PCRs, Fig.S5a,b,c). Amplicon sequencing data also revealed that different *Stenotrophomonas* species/strains (OTUs) colonized the rhizosphere and roots, which is in line with the multiple strains obtained by the culture-dependent method. Among the four strains, only the 16S rRNA of *Stenotrophomonas* spp. R80 (hereafter referred as SR80) had 100% nucleotide similarity match with the sequences of OTU 89589 and OTU 41442 (Fig.S5a,b,c). The whole genome of SR80 was then sequenced and analyzed (Fig.S6&7), through which the 16S rRNA region was obtained, which was 1,532 bp in length and had 100% sequence similarity to OTU 89589 and OTU 41442 (Fig.S6a,b). This suggests that SR80 was the bacterium enriched in the wheat microbiome by *Fp*-infection (Fig.3a,b,c,d, Fig.S6a,b). The SR80 genome had the highest percent of Average Nucleotide Identity (ANI) (97.71%) to a *Stenotrophomonas rhizophila* strain QL-P4 that was isolated from a pine tree leaf, suggesting it is a member of *S.rhizophila* species but possibly a different strain.

**Fig.3.**
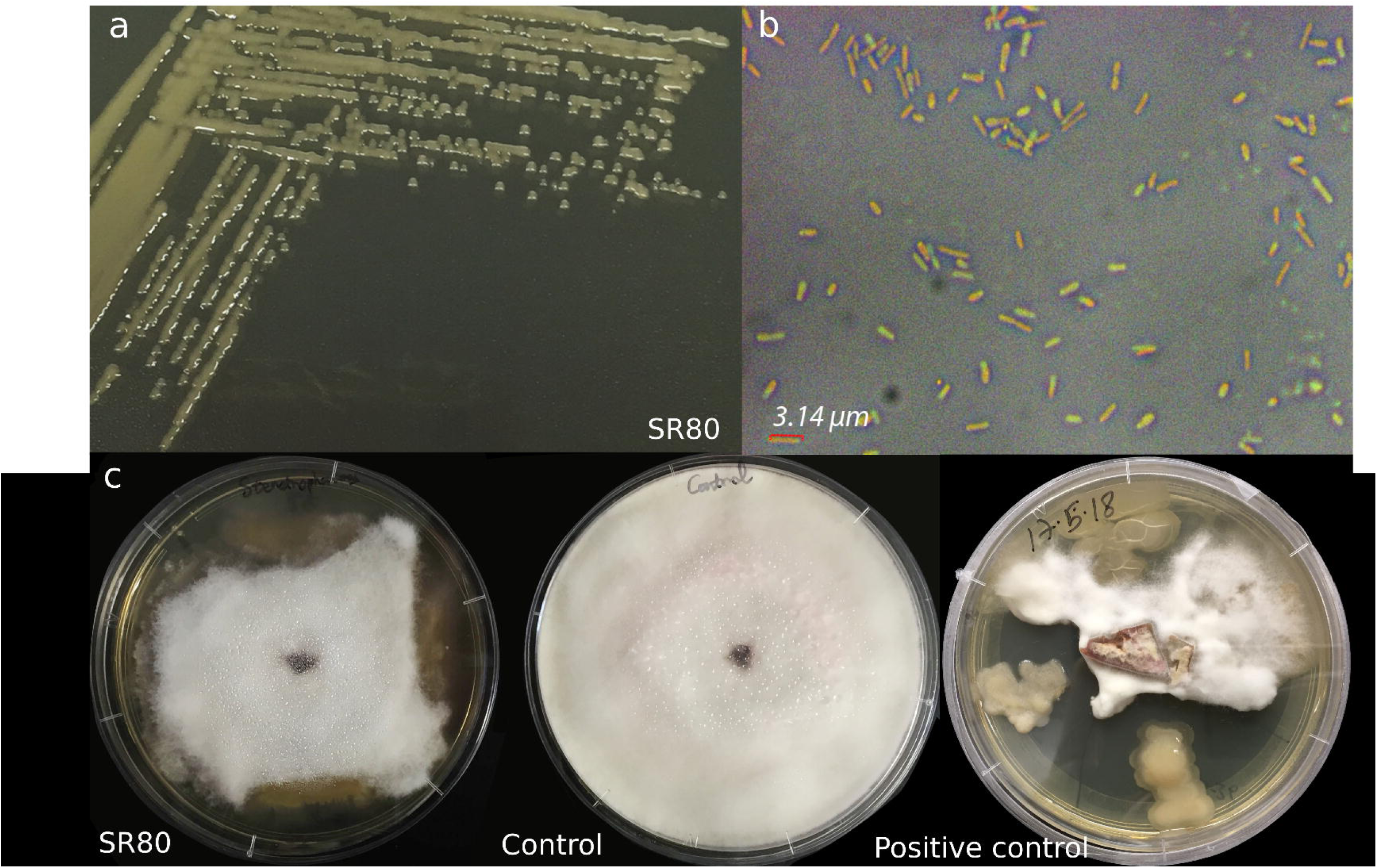
Morphology of SR80 isolated from wheat rhizosphere soil and root endosphere. Colony morphology of SR80 grown for 24 h on trypticase soy agar (TSA) (a). Bacterial cell morphology under a light microscope (b). Slight inhibition of the bacterium on *Fp* growth on a dual media (1/2 PDA+1/2 TSA) (c). Bacteria used as positive controls at the bottom right were two strains of *Burkholderia* spp.

The predicted genome size of SR80 is 4,234,260 bp containing 3,720 detected genes (Fig.S7). Class of genes (COG) annotation was performed on the whole genome and 3,052 protein sequences were successfully annotated. Known virulence factors and plant cell wall degrading enzymes were absent, but genes encoding proteins involved in symbiosis, host-cell interaction and activation of plant defense signaling pathways were detected. Genes coding for bacterial flagellin (*flg22*), chitin (*AvrXa21*), and Type I∼ Type IV secretion systems were detected (Fig.S8), as were genes *fliR* and *flhB* which code for effector proteins that interact with the plant immune system and induce the plant’s primary responses. Genes encoding proteins involved in biofilm formation, chemotaxis, lipopolysaccharide (LPS) biosynthesis, the quorum sensing system and reactive oxygen species (ROS) scavenging (e.g., superoxide dismutase) are also present in multiple copies. These indicate that (i) SR80 can cope with plant immune responses that may prevent bacterial colonization, and (ii) this bacterium could be well adapted to the root environment. SR80 was further studied in antifungal tests against four virulent *Fp* strains (including the *Fp* used in the field experiment) on dual media, but no noticeable inhibition was observed (Fig.3c).

### SR80 promoted plant growth and defense

To evaluate whether SR80 affects wheat growth and defense, we treated the *Fp*-infected variety of durum wheat (Jandaroi) with SR80 in glasshouse experiments (Fig.4). Overall, inoculation with SR80 at the time of sowing significantly promoted plant growth and conferred protection against *Fp* compared with the non-inoculated control (Fig.4a,b,c,d). Within 15 days, plant biomass in both, below-ground (roots, +156%) and above-ground (shoots, +124%), compartments substantially increased by SR80 inoculation (Fig.4b). This is consistent with the observation that the diameter of the stem base was significantly larger at the two time points monitored, although the height of the SR80-inoculated plants was not larger than the control within the timeframe tested (Fig.4c). Moreover, plant survival rates were significantly higher in the SR80-inoculated group than the untreated group when subjected to *Fp* infections. The plants in the untreated control all died from *Fp* infection within 15 days (Fig.4d). qPCR quantification did not reveal a decrease in *Fp* load after the bacterial inoculation (Fig.4f), which is consistent with our plate assay results. The bacterial communities in soils were profiled at three time points (5, 10 and 15 days). We found that the abundance of the added SR80 in soils gradually decreased to a stable level 10 days after seed germination (∼2.4×10^7^ cells g^-1^ soil) (Fig.4e), in agreement with the relative abundance of SR80 in the soil microbiome continuously declining within the timeframe, from 70.0% to 40.74% (Fig.S9). The alpha diversity of soil microbial communities gradually increased in all groups within 15 days, with the highest diversity observed in the uninoculated control (Fig.S10).

**Fig.4.**
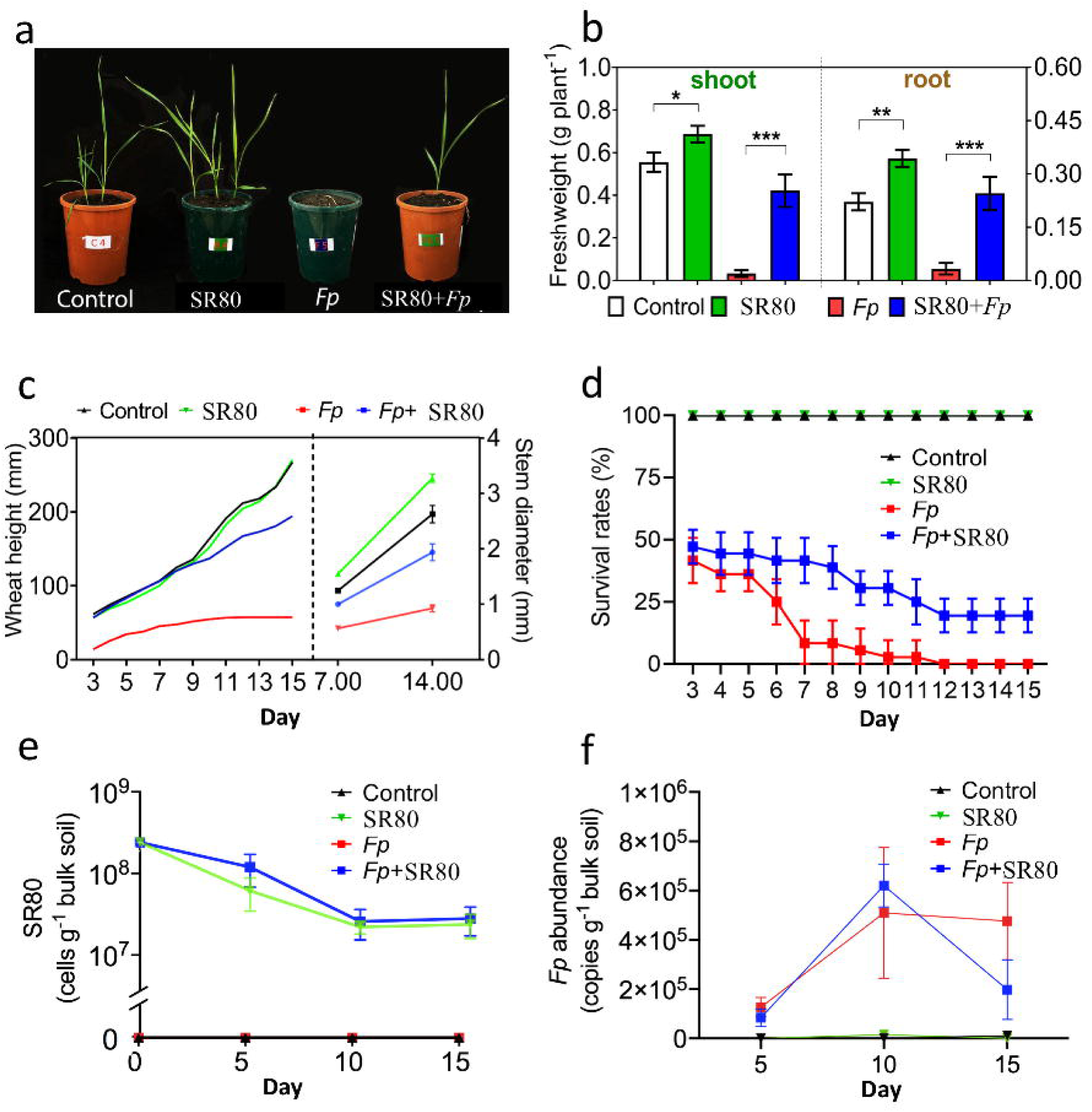
Effects of SR80 inoculation on wheat growth and survival. Wheat phenotypes 14 days post treatment applications (a). Control: no inoculation, SR80: inoculation with SR80 only, *Fp*: inoculation with *Fp* only, SR80+*Fp*: dual inoculation with SR80 and *Fp*. Biomass accumulation of wheat shoots and roots 15 days after the treatment applications (b). Increase in plant height and stem size over 15 days from the application of treatments (c). Wheat survival rates under different treatments (d). Changes in abundance of SR80 (e) and *Fp* (f) abundance in soils. Asterisks indicate significant differences between treatments (\**P*□<□0.05, \*\**P*□<□0.01, \*\*\**P*□<□0.001). Error bars in (b)-(f) represent standard errors of the mean (n=6).

### SR80 manipulated plant defense signaling pathways

SR80 did not significantly inhibit *Fp* in soil or on agar plates, indicating that a direct inhibition of the pathogen is not the mechanism for this bacterium to benefit wheat survival and growth. We hypothesized that SR80 modulates expression of genes involved in plant defense. An additional glasshouse experiment was conducted to test this hypothesis, in which plants were inoculated with both the pathogen *Fp* and the beneficial bacterium SR80 and evaluated for expression of marker genes associated with plant defense signaling pathways. We found that when SR80 is present in soils, JA and SA signaling pathways were significantly activated in wheat but only in the presence of *Fp*. Expression of genes, including *TaAOS* (+3.1 fold) (Fig.5a), *TaNPR1* (+2.2 fold) (Fig.5b), *Lipase* (+6.0 fold) (Fig.5c), *WRKY78* (+3.5 fold) (Fig.5d), *PR2* (+3.0 fold) (Fig.5f), and *PR3* (+2.3 fold) (Fig.5g), was enhanced by SR80 treatments (Fig.5g). In contrast, expression of *PR4* marginally decreased by the bacterial treatment without *Fp* infection. Other PR genes also showed a decrease although were not statistically significant (Fig.5h).

**Fig.5.**
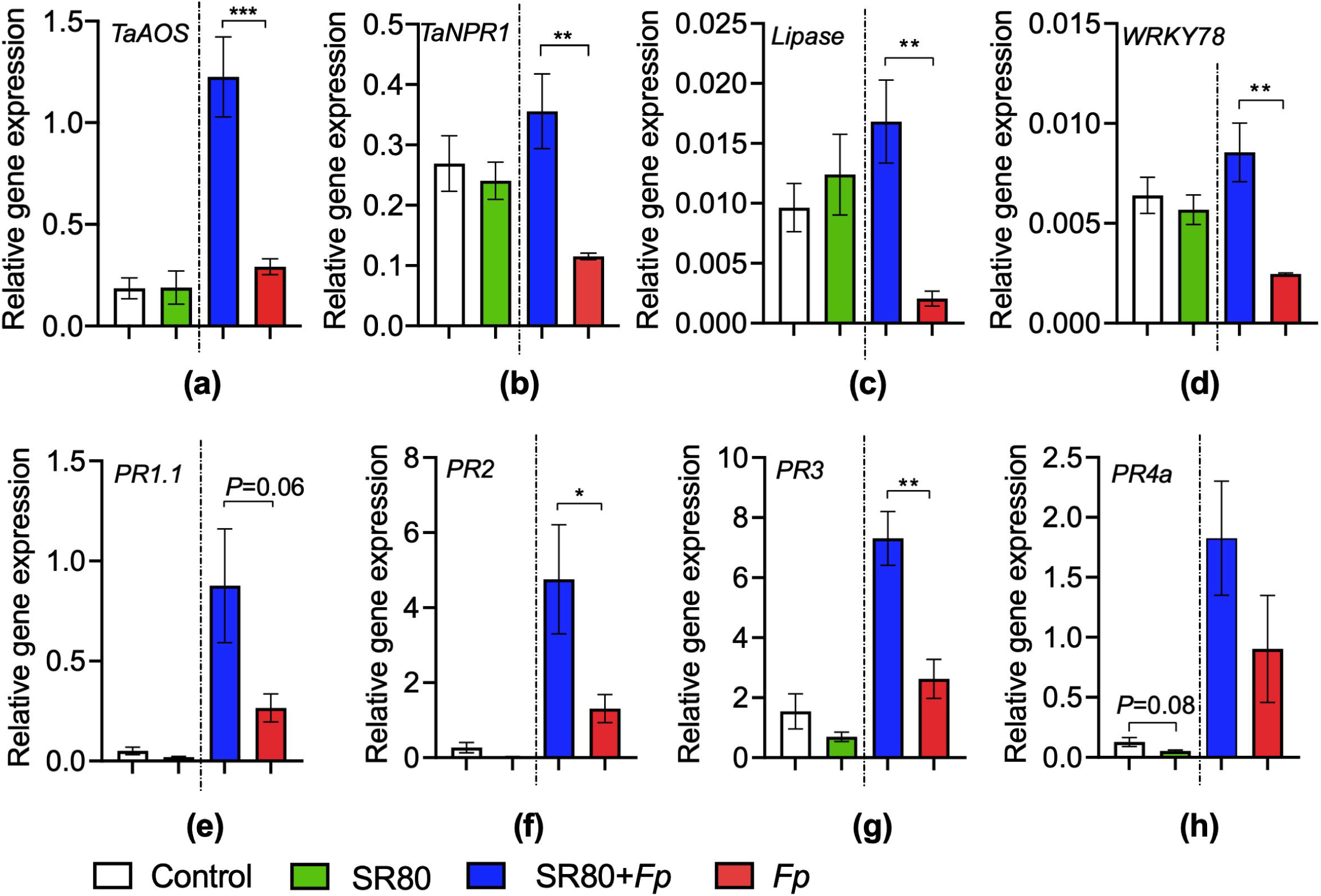
Effects of SR80 and *Fp* inoculations on the transcription of genes associated with JA and SA signaling pathways. Glasshouse experiment 1 (left) and 2 (right) were separated by black dash lines on each graph. Asterisks indicate significant differences between treatments (\**P*□<□0.05, \*\**P*□<□0.01, \*\*\**P*□<□0.001). Error bars represent standard errors of the means (n =□6).

**Fig.6.**
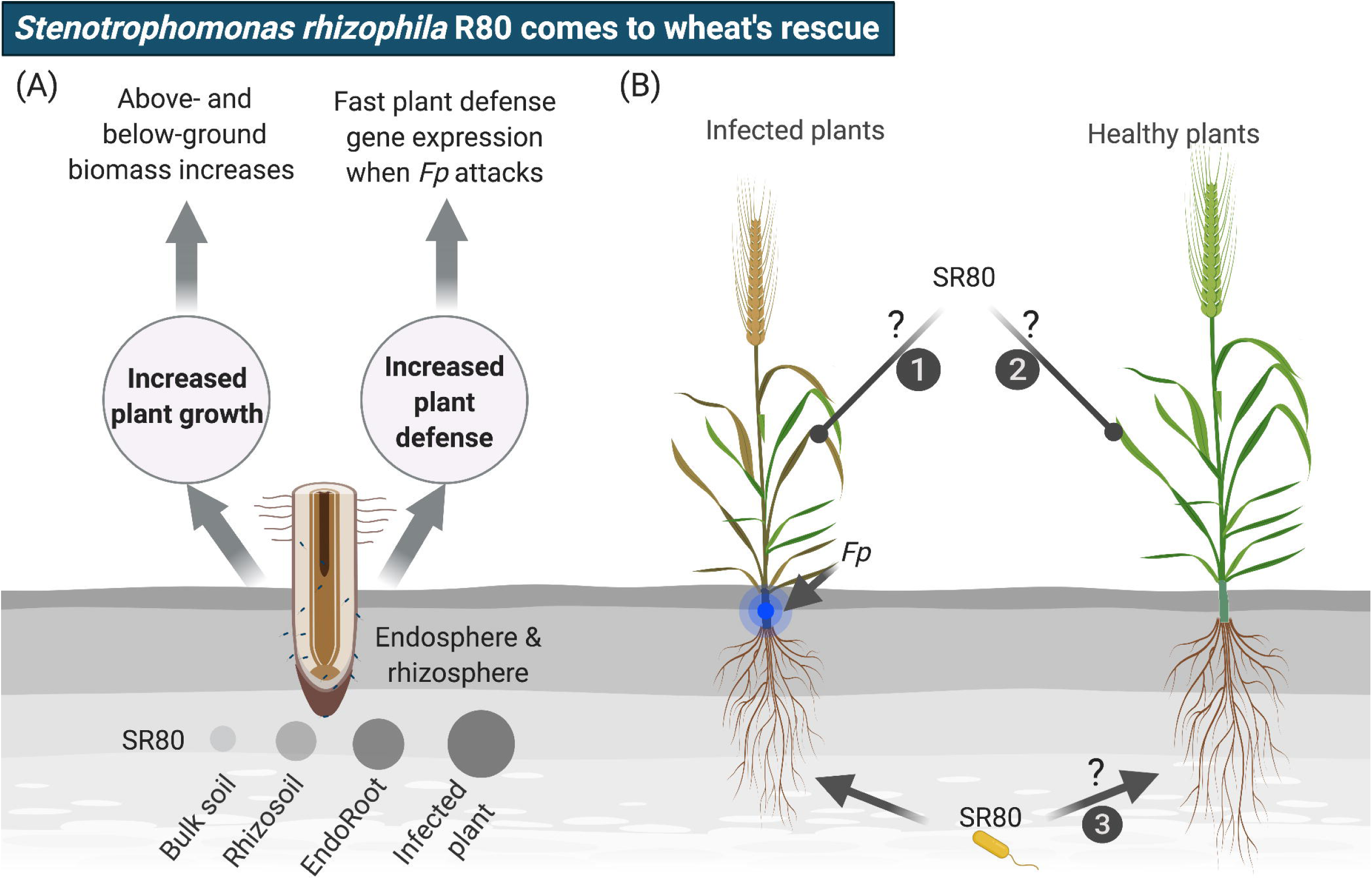
Schematic drawing illustrating biological implications of SR80 on plant growth and defense. The bacterium SR80 can modulate wheat growth and defense, which seems to be mediated by the plant immune system. The plant favors the enrichment of the bacterium from the rhizosphere soils and to the root endosphere. Moreover, when exposed to *Fp*, the wheat plant favors the enrichment with even more SR80 in the rhizosphere. Such an active recruitment process is accompanied by an enhanced defense gene expression in leaves. Future research can investigate (i) whether plant leaves enrich the bacterium (process 1), and (ii) whether allelopathic effects lead to the enrichment of the bacterium in the neighboring threatened plants (process 2&3).

## Discussion

Our study demonstrates that CR disease in wheat plants leads to an enrichment of the beneficial bacterium SR80 in the rhizosphere soil and root endosphere. The assembled SR80 was able to significantly enhance plant growth at both the above- and below-ground and induce resistance against the CR disease in glasshouse experiments. Despite strains of *S.rhizophila* having been shown to suppress mycelial growth in various fungi at times through the production of volatiles (Kai *et al.*, 2007), SR80 did not directly inhibit *Fp* growth on plates or in soil. However, SR80 can upregulate plant defense signalings (e.g. JA and SA) in shoots with the presence of the pathogen. These findings provide a novel mechanism of tripartite interactions between the devastating fungal pathogen, the plant host and plant-associated microbiota, and revealed a bacterium that may have acted as an early warning system for the onset of plant defense.

### SR80 intimately interacts with plant immune systems and mediates a disease resistance

SR80 inoculation tends to suppress the expression of PR-related defense genes in wheat shoots without *Fp* presence in soils. Consistent with this finding, suppression of the plant root immunity by non-pathogenic *Pseudomonas* species has recently been observed in *Arabidopsis thaliana*, which may have facilitated the bacterial colonization on roots (Yu *et al.*, 2019). These suggest that quenching plant immune responses is an emerging strategy for plant-associated bacteria to prevent constitutive activation of plant immune responses. Rhizosphere and endosphere microbiota are rich in microbe-associated molecular patterns (MAMPs), which can trigger the first layer of plant immune defense that restricts pathogen reproduction, MAMP-triggered immunity. SR80 undoubtedly possesses a range of components acting as MAMPs (e.g. flagellin), but how the bacterium avoids eliciting strong immune responses in plants is unclear. Recent conceptual and experimental framework indicates that plant-associated commensal microbes can alleviate/avoid plant immune responses by (i) producing enzymes to scavenge reactive oxygen species (ROS) production of the plant (Sessitsch *et al.*, 2012), (ii) producing organic acids to quench local plant immune responses (Yu *et al.*, 2019), and (iii) modifying, degrading or changing the structure of MAMPs (Liu *et al.*, 2020). Our whole genome annotation data suggest that SR80 contains multiple genes encoding key components that are functioning by these mechanisms.

Interestingly, upon challenge with *Fp*, pre-colonization by SR80 on wheat seedings induced a much stronger plant defense that increased plant survival rates relative to untreated plants. This suggests that SR80 possesses elicitors that triggered Induced Systemic Resistance (ISR) and ‘primed’ the plant for faster and more pronounced responses to pathogen attack. Both the SA and JA signaling pathways of the plant were activated, which is quite unusual because SA and JA signaling pathways are mostly antagonistic, although non-canonical mechanisms of synergism have been reported (Thaler *et al.*, 2012). In monocots, the relationships between the disease signaling pathways and the nutrient acquisition behavior of the pathogen is less understood.

*Fp* behaves as a hemi-biotroph as there is an extended initial phase that plants show no symptoms, after which necrotic lesions develop. Two oxalotrophic strains of the same genus as SR80, *Stenotrophomonas*, were isolated from the tomato rhizosphere and have been shown to upregulate *PR-1*, a marker gene for the SA pathway (Marina *et al.*, 2019). However, these strains did not influence the expression of *PDF1.2*, a marker gene for JA/ethylene (ET) signaling pathway (Marina *et al.*, 2019). In line with our findings, a strain of plant growth promoting *Stenotrophomonas maltophilia*, isolated from the rhizosphere of sorghum, increased synthesis of a range of enzymes associated with defense against fungal pathogens when inoculated on wheat, including peroxidase, β-1,3 glucanase, polyphenol oxidase and phenylalanine ammonia lyase (Singh & Jha, 2017). Overall, our findings suggest that SR80 can play a key role in the three-way interactions with the host plant immunity and the fungal pathogen via a mechanism that enhances the plant defence and growth.

### SR80 was enriched in root environments upon Fp infection

In this study, we used amplicon sequencing and qPCR for the analysis of plant microbiomes, which can detect differences in microbial community structure and estimate the abundance of microbial taxa in soils. Our results revealed that SR80 was thriving at high abundances around roots of the *Fp*-infected plants (up to 3.7×10^7^ cells g^-1^ rhizosphere soils). *S. maltophilia* was also present at high titers in root samples of oilseed rape, about 1.1×10^7^ colony forming units (cfu) g^-1^ wet weight root (Berg *et al.*, 1996). A minimum of 10^5^ cfu g^-1^ root has been reported to be required for the onset of ISR by certain beneficial microbes (Pieterse *et al.*, 2014). Accordingly, the quantity of SR80 accumulated in the rhizosphere and roots of the infected wheat is likely have reached sufficient numbers to induce plant defense responses and confer a protective barrier to restrict *Fp*. This is consistent with upregulated defenses in wheat plants under both the glasshouse and field conditions by the presence of SR80. However, we noticed that the developed CR symptoms persisted throughout the life cycle of the plant in the field and glasshouse experiments, which suggests that the recruitment of SR80 unlikely cures an existing infection, but may instead increase the plant tolerance to the disease.

Recent studies in dicot plants including *A.thaliana* and pepper (*Capsicum annuum* L. cv. Bukwang) are in line with our findings as attack of foliar tissues by pathogens or insect herbivores resulted in plant-mediated changes in the rhizosphere microbial communities (Kong *et al.*, 2016; Berendsen *et al.*, 2018). Particular microbial taxa affiliated to *Stenotrophomonas* spp. were similarly enriched in the diseased plants, which suggests that this genus is commonly influenced by different plant biotic stresses. *Stenotrophomonas* spp. including *S. rhizophila* possess a strong capacity to colonize root tissues (Hayward *et al.*, 2010), which allows the bacterium to intimately interact with the host plant. Under the conditions tested in our study, wheat preferentially selected certain microbial phylotypes (e.g. Gammaproteobacteria) including SR80, resulting in a continuous increase of SR80 abundance in the wheat microbiome as habitats shifted from bulk soil to the root endosphere.

In fact, there is evidence to suggest that healthy plants assemble different microbial populations to assist their growth and development (Yuan *et al.*, 2015; Qiao *et al.*, 2017; Imchen *et al.*, 2019). CR disease also modulated the wheat microbiome towards accumulation of more SR80 in the rhizospheric and endophytic microbial communities, suggesting that disease infection can be a critical driver for the plant microbiome assembly, and a cry for help strategy can be triggered to allow plants to actively seek help from soils to cope with biotic stresses.

It is worth pointing out that not all plant diseases induce microbial changes in the plant microbiome. For example, infection by the necrotrophic pathogen *Botrytis cinerea* in *A.thaliana* did not clearly promote microbes in the rhizosphere although it did activate the plant JA-dependent defense pathway (Berendsen *et al.*, 2018). This indicates the pattern of disease-induced changes in the plant microbiome is defined by the particular interactions occurring between a specific plant species and a pathogen. Furthermore, enrichment of particular microbes in plant microbiomes can at times promote the fungal spore germination and virulence (Partida-Martinez & Hertweck, 2005; Seneviratne *et al.*, 2007). *Burkholderia* spp. for example, forms a symbiotic relationship with the pathogenic fungus *Rhizopus*, the causative agent of rice seedling blight (Partida-Martinez & Hertweck, 2005). This bacterium produces a toxin that is required for the pathogen to colonize its rice host. In this case, the enrichment of a bacterium helps in the establishment of a fungal disease.

Collectively, our findings suggest that infection of wheat plants by CR disease results in the recruitment of a beneficial rhizospheric microbe that has the potential to help plant growth and protect plants via modulation of plant defense capacity. The enriched microbe acts as an early warning agent, rapidly activating the JA and SA signaling pathways upon pathogen invasion in plants. This work advances the current understanding of plant-microbe interaction research and supports the coevolution theory of mutualism between the plant and microbes.

## Acknowledgements

The authors would like to acknowledge the financial support from the Australian Research Council (DP1094749; DP170104634; DP190103714). We thank Dr Friday Obanor for providing us *Fp* strains and Dr Anna Balzer for helping with plant sampling and soil collection. We also thank Associate Professor Mark Dieters for providing wheat seeds and Dr Laura E. Brettell for proofreading this article.

## Author contributions

H.L., P.S., L.C.C conceptualized the idea, which was further refined by B.K.S; H.L., L.C.C., and C.P. did the plant and soil sampling; H.L. and J.L. processed the plant and soil samples and did the measurements for glasshouse treatments; H.L. analyzed the data and wrote the first draft with significant inputs from B.K.S; all authors have critically read and revised the manuscript.

## Data availability

Data of this study are available at 10.6084/m9.figshare.12464465. Other relevant data are available upon request.

## References

Akinsanmi O, Mitter V, Simpfendorfer S, Backhouse D, Chakraborty S. 2004. Identity and pathogenicity of Fusarium spp. isolated from wheat fields in Queensland and northern New South Wales. Crop Pasture Sci 55(1): 97–107.

Álvarez-Pérez S, Lievens B, Fukami T. 2019. Yeast & bacterium interactions: the next frontier in nectar research. Trends Plant Sci 24(5): 393–401.

Berendsen RL, Vismans G, Yu K, Song Y, de Jonge R, Burgman WP, Burmølle M, Herschend J, Bakker PAHM, Pieterse CMJ. 2018. Disease-induced assemblage of a plant-beneficial bacterial consortium. ISME J 12(6): 1496–1507.

Berg G, Marten P, Ballin G. 1996. Stenotrophomonas maltophilia in the rhizosphere of oilseed rape — occurrence, characterization and interaction with phytopathogenic fungi. Microbiol Res 151(1): 19–27.

Bollet C, Davin-Regli A, De Micco P. 1995. A simple method for selective isolation of Stenotrophomonas maltophilia from environmental samples. Appl Environ Microbiol 61(4): 1653–1654.

Chakraborty S, Newton AC. 2011. Climate change, plant diseases and food security: an overview. Plant Pathol 60(1): 2–14.

Chakraborty S, Obanor F, Westecott R, Abeywickrama K. 2010. Wheat crown rot pathogens Fusarium graminearum and F. pseudograminearum lack specialization. Phytopathology 100(10): 1057–1065.

Dixon P. 2003. VEGAN, a package of R functions for community ecology. J Veg Sci 14(6): 927–930.

Durán P, Thiergart T, Garrido-Oter R, Agler M, Kemen E, Schulze-Lefert P, Hacquard S. 2018. Microbial interkingdom interactions in roots promote Arabidopsis survival. Cell 175(4): 973-983.e914.

Edwards J, Johnson C, Santos-Medellín C, Lurie E, Podishetty NK, Bhatnagar S, Eisen JA, Sundaresan V. 2015. Structure, variation, and assembly of the root-associated microbiomes of rice. Proc Natl Acad Sci USA 112(8): E911–E920.

Engelbrektson A, Kunin V, Wrighton KC, Zvenigorodsky N, Chen F, Ochman H, Hugenholtz P. 2010. Experimental factors affecting PCR-based estimates of microbial species richness and evenness. ISME J 4(5): 642.

Glass NL, Donaldson GC. 1995. Development of primer sets designed for use with the PCR to amplify conserved genes from filamentous ascomycetes. Appl Environ Microbiol 61(4): 1323–1330.

Grady KL, Sorensen JW, Stopnisek N, Guittar J, Shade A. 2019. Assembly and seasonality of core phyllosphere microbiota on perennial biofuel crops. Nat Commun 10(1): 4135.

Hayward AC, Fegan N, Fegan M, Stirling GR. 2010. Stenotrophomonas and Lysobacter: ubiquitous plant-associated gamma-proteobacteria of developing significance in applied microbiology. J Appl Microbiol 108(3): 756–770.

Ihrmark K, Bödeker I, Cruz-Martinez K, Friberg H, Kubartova A, Schenck J, Strid Y, Stenlid J, Brandström-Durling M, Clemmensen KE. 2012. New primers to amplify the fungal ITS2 region–evaluation by 454-sequencing of artificial and natural communities. FEMS Microbiol Ecol 82(3): 666–677.

Imchen M, Kumavath R, Vaz ABM, Góes-Neto A, Barh D, Ghosh P, Kozyrovska N, Podolich O, Azevedo V. 2019. 16S rRNA gene amplicon based metagenomic signatures of rhizobiome community in rice field during various growth stages. Front Microbiol 10(2103).

Kai M, Effmert U, Berg G, Piechulla B. 2007. Volatiles of bacterial antagonists inhibit mycelial growth of the plant pathogen Rhizoctonia solani. Arch Microbiol 187(5): 351–360.

Kong HG, Kim BK, Song GC, Lee S, Ryu C-M. 2016. Aboveground whitefly infestation-mediated reshaping of the root microbiota. Front Microbiol 7(1314).

Kwak M-J, Kong HG, Choi K, Kwon S-K, Song JY, Lee J, Lee PA, Choi SY, Seo M, Lee HJ, et al. 2018. Rhizosphere microbiome structure alters to enable wilt resistance in tomato. Nat Biotechnol 36(11): 1100–1109.

Lagesen K, Hallin P, Rødland EA, Stærfeldt H-H, Rognes T, Ussery DW. 2007. RNAmmer: consistent and rapid annotation of ribosomal RNA genes. Nucleic Acids Res 35(9): 3100–3108.

Liu H, Brettell LE. 2019. Plant defense by VOC-induced microbial priming. Trends Plant Sci 24(3): 187–189.

Liu H, Brettell LE, Qiu Z, Singh BK. 2020. Microbiome-mediated stress resistance in plants. Trends Plant Sci 25(8), 733–743.

Liu H, Carvalhais LC, Kazan K, Schenk PM. 2016. Development of marker genes for jasmonic acid signaling in shoots and roots of wheat. Plant Sig Behav 11(5): e1176654.

Liu H, Macdonald CA, Cook J, Anderson IC, Singh BK. 2019. An ecological loop: host microbiomes across multitrophic interactions. Trends Ecol Evol 34(12): 1118–1130.

Marina M, Romero FM, Villarreal NM, Medina AJ, Gárriz A, Rossi FR, Martinez GA, Pieckenstain FL. 2019. Mechanisms of plant protection against two oxalate-producing fungal pathogens by oxalotrophic strains of Stenotrophomonas spp. Plant Mol Biol 100(6): 659–674.

Melloy P, Hollaway G, Luck J, Norton R, Aitken E, Chakraborty S. 2010. Production and fitness of Fusarium pseudograminearum inoculum at elevated carbon dioxide in FACE. Global Change Biol 16(12): 3363–3373.

Mendes LW, Raaijmakers JM, de Hollander M, Mendes R, Tsai SM. 2018. Influence of resistance breeding in common bean on rhizosphere microbiome composition and function. ISME J 12(1): 212–224.

Nurk S, Meleshko D, Korobeynikov A, Pevzner PAJGr. 2017. metaSPAdes: a new versatile metagenomic assembler. 27(5): 824–834.

Partida-Martinez LP, Hertweck C. 2005. Pathogenic fungus harbours endosymbiotic bacteria for toxin production. Nature 437(7060): 884–888.

Peiffer JA, Spor A, Koren O, Jin Z, Tringe SG, Dangl JL, Buckler ES, Ley RE. 2013. Diversity and heritability of the maize rhizosphere microbiome under field conditions. Proc Natl Acad Sci USA 110(16): 6548–6553.

Percy CD, Wildermuth GB, Sutherland MW. 2012. Symptom development proceeds at different rates in susceptible and partially resistant cereal seedlings infected with Fusarium pseudograminearum. Australas Plant Path 41(6): 621–631.

Pieterse CMJ, Zamioudis C, Berendsen RL, Weller DM, Wees SCMV, Bakker PAHM. 2014. Induced systemic resistance by beneficial microbes. Annu Rev Phytopathol 52(1): 347–375.

Qiao Q, Wang F, Zhang J, Chen Y, Zhang C, Liu G, Zhang H, Ma C, Zhang J. 2017. The variation in the rhizosphere microbiome of cotton with soil type, genotype and developmental stage. Sci Rep 7(1): 3940.

Ramakers C, Ruijter JM, Deprez RHL, Moorman AFM. 2003. Assumption-free analysis of quantitative real-time polymerase chain reaction (PCR) data. Neurosci Lett 339(1): 62–66.

Seneviratne G, Zavahir JS, Bandara WMMS, Weerasekara MLMAW. 2007. Fungal-bacterial biofilms: their development for novel biotechnological applications. World J Microbiol Biotechnol 24(6): 739.

Sessitsch A, Hardoim P, Döring J, Weilharter A, Krause A, Woyke T, Mitter B, Hauberg-Lotte L, Friedrich F, Rahalkar M. 2012. Functional characteristics of an endophyte community colonizing rice roots as revealed by metagenomic analysis. Mol Plant Microbe Interact 25(1): 28–36.

Singh RP, Jha PN. 2017. The PGPR Stenotrophomonas maltophilia SBP-9 augments resistance against biotic and abiotic stress in wheat plants. Front Microbiol 8: 1945.

Thaler JS, Humphrey PT, Whiteman NK. 2012. Evolution of jasmonate and salicylate signal crosstalk. Trends Plant Sci 17(5): 260–270.

Wildermuth GB, McNamara RB. 1994. Testing wheat seedlings for resistance to crown rot caused by Fusarium graminearum group 1. Plant disease 78(10): 949–953.

Yu K, Liu Y, Tichelaar R, Savant N, Lagendijk E, van Kuijk SJ, Stringlis IA, van Dijken AJ, Pieterse CM, Bakker PA. 2019. Rhizosphere-associated Pseudomonas suppress local root immune responses by Gluconic Acid-mediated lowering of environmental pH. Curr Biol 29(22): 3913-3920. e3914.

Yuan J, Chaparro JM, Manter DK, Zhang R, Vivanco JM, Shen Q. 2015. Roots from distinct plant developmental stages are capable of rapidly selecting their own microbiome without the influence of environmental and soil edaphic factors. Soil Biol Biochem 89: 206–209.

