## supplementary materials for "Evidence for the plant recruitment of beneficial microbes to suppress soil-borne pathogen"

*^1^Hawkesbury Institute for the Environment, Western Sydney University, Penrith, NSW 2753, Australia; ^2^School of Agriculture and Food Sciences, The University of Queensland, Saint Lucia, Queensland 4072, Australia; ^3^Centre for Horticultural Science, Queensland Alliance for Agriculture and Food Innovation, The University of Queensland, Saint Lucia, Queensland 4102 Australia; ^4^Centre for crop health, University of Southern Queensland, Toowoomba, Queensland 4350, Australia; ^5^Institute of Environment and Sustainable Development, Banaras Hindu University, Varanasi-221005, Uttar Pradesh, India; ^6^Global Centre for Land-Based Innovation, Western Sydney University, Penrith, NSW 2753, Australia. ^7^These authors contributed equally.*

**Supplementary methods**

**Standard PCR:** PCRs were performed in a 25 µL reaction volume containing: 14.75 µL of ultra-pure water, 5 µL of 5×Phire buffer (Thermo Scientific), 1.25 µL of dNTPs (10 µM), 1.25 µL of a 10 µM forward primer, 1.25 µL of a 10 µM reverse primer, 0.5 µL of Phire^®^ Hot Start II (Thermo Scientific), and 1 µL of DNA template (10- 20 ng DNA). PCR thermal conditions were 30 s at 98°C for initial denaturation, 30 cycles of 15 s at 98°C, 30 s for annealing at specific temperatures for each primer pair (Table S2) and 45 s at 72°C for elongation; followed by 7 min at 72°C for the final extension.

**qPCR and qRT-PCR:** plant and soil DNA samples were normalized to 10 ng µL^-1^ and 5 ng µL^-1^, respectively, for qPCR/qRT-PCR analysis. The PCRs were performed in a 10 µL reaction volume containing 5 µL of SYBR Green PCR master mix, 1 µL of 3 µM mix of forward and reverse primers and 4 µL DNA (5~10 ng for gDNA, ~17 ng for cDNA). Cycling conditions included 40 cycles of a denaturation step at 95°C for 15 s and an annealing/extension step at specific temperatures for each primer pair for 1 min (Table S2), followed by generation of a melt curve analysis by heating to 95°C for 2 min, followed by 60 °C for 15 s, and 95 °C for 15 s.

**Bioinformatics**

*Bacterial and archaeal community analyses*

Raw sequence data were processed as previously described (Liu *et al.*, 2016; Liu *et al.*, 2017; Liu *et al.*, 2019). Briefly, primer sequences were removed from each FASTQ file and the header line of each sequence was then modified to contain a sample ID using QIIME v1.9.1. Sequences were then quality-filtered using the QIIME script split_libraries.py with the homopolymer filter deactivated (Caporaso et al. 2010). The forward reads from each sample were concatenated into a single file and checked for chimeras against GreenGenes database (version October, 2013) using UCHIME ver. 3.0.617 (Edgar et al. 2011). Homopolymer errors were corrected using Acacia (Bragg et al. 2012). The resulting sequences were processed by the following procedures using QIIME: (i) sequences were clustered at 97% similarity using UCLUST, (ii) GreenGenes taxonomy was assigned to the cluster representatives using BLAST, and (iii) tables with the abundance of different Operational Taxonomic Unit (OTUs) and their taxonomic assignments in each sample were generated. The number of reads was rarefied to 3,600 and 2,900 sequences per sample for the rhizosphere soil and root samples, respectively, by re-sampling the OTU table. Richness and evenness of plant and soil microbiomes were calculated using QIIME as Simpson’s diversity, observed (Sobs) and predicted OTU abundance (Chao1).

*Fungal community analyses*

The obtained FASTQ files from ITS sequencing were processed using QIIME2 (version 2019.7; http://qiime2.org/) (Bolyen *et al.*, 2019). Briefly, the quality of FASTQ files was assessed by FastQC (Andrews, 2010). DADA2 (Callahan *et al.*, 2016) was used in QIIME2 for removal of primer sequences, error-correction, quality filtering, chimera removal and sequence variant calling. Forward and reverse sequences were merged after trimming off primers and truncating sequences at 260 bp (forward) and 240 bp (reverse) to keep only high quality sequences (Q>20). The feature sequences (OTUs) obtained via this step were summarized and assigned with taxonomy information using an RDP classifier (Cole *et al.*, 2008) pre-trained using UNITE database (released 04/07/2014). The number of reads was rarefied to 10,000 per sample by re-sampling the OTU table. The mean number of observed (Sobs) OTUs, Simpson’s and Shannon diversity index values were calculated within QIIME2. The 16S rRNA and ITS amplicon sequences associated with this study have been deposited in the NCBI SRA under accession code PRJNA436828.

**Table S1** Soil chemical characteristics.

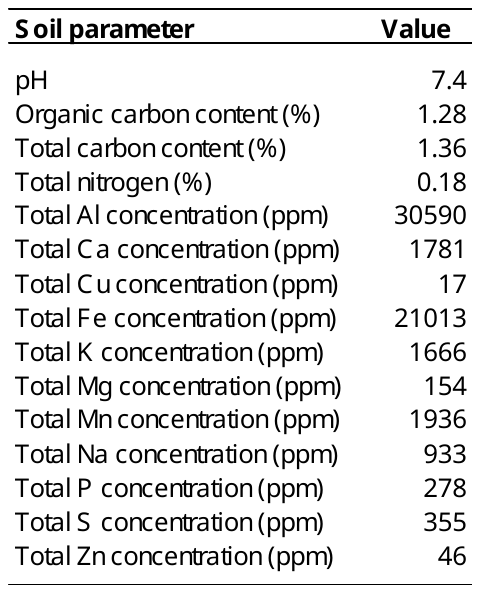

Total nitrogen was measured using the method from the handbook section: 6B2, p75 (Rayment and Lyons, 2011). Mineral elements were measured as described by the USEPA method 3052, entitled “Microwave assisted acid digestion of siliceous and organically based matrices”, Kingston HM and Walter PJ (Rayment & Lyons, 2011).

**Table S2** Management history of the experimental field at Wellcamp. During the fallow, paddocks were sprayed with herbicide and cultivated as needed for weed control.

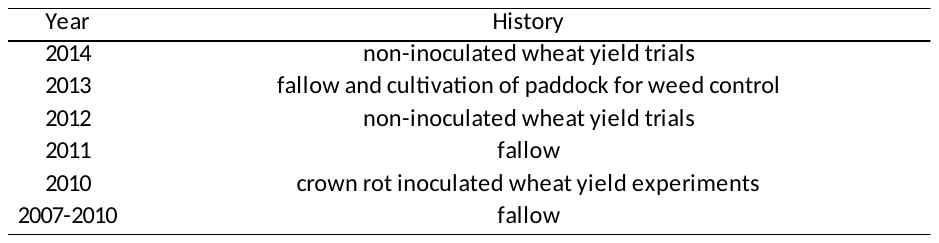

**Table S3** Primer sequences used in this study.

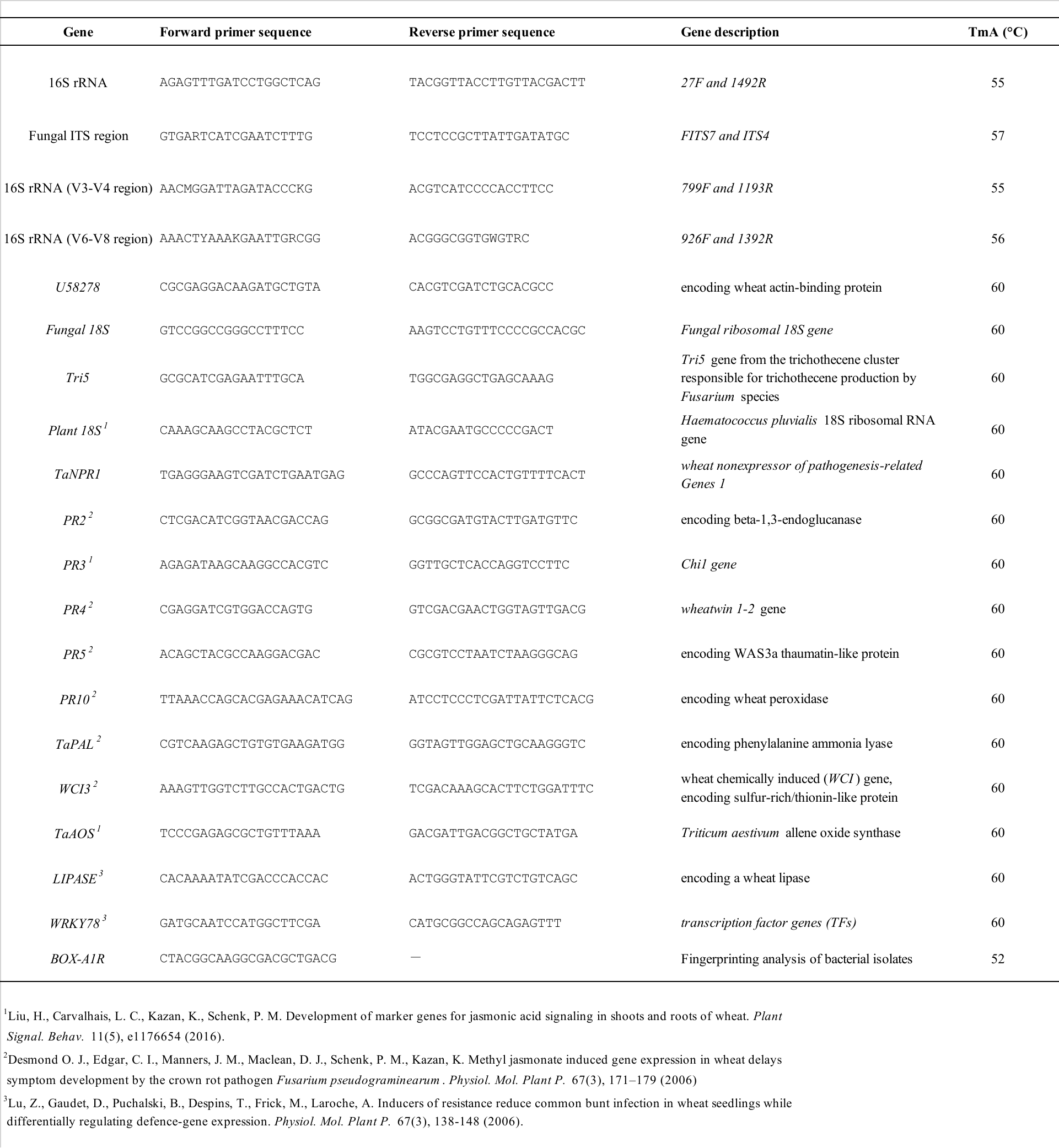

**Table S4** Disease-induced effects on alpha diversity of plant and soil microbial communities. The values were based on 3,600 (rhizosphere soil) and 2,900 (root endosphere) rarefied sequences per sample. Errors are SEs (n=40 healthy, n=18 infected). Distinct lowercase letters show significant differences between healthy and diseased soil/root samples.

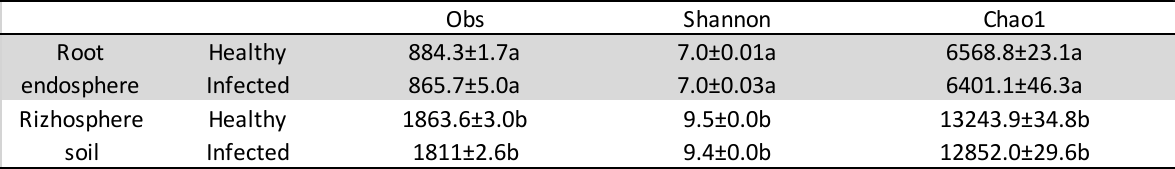

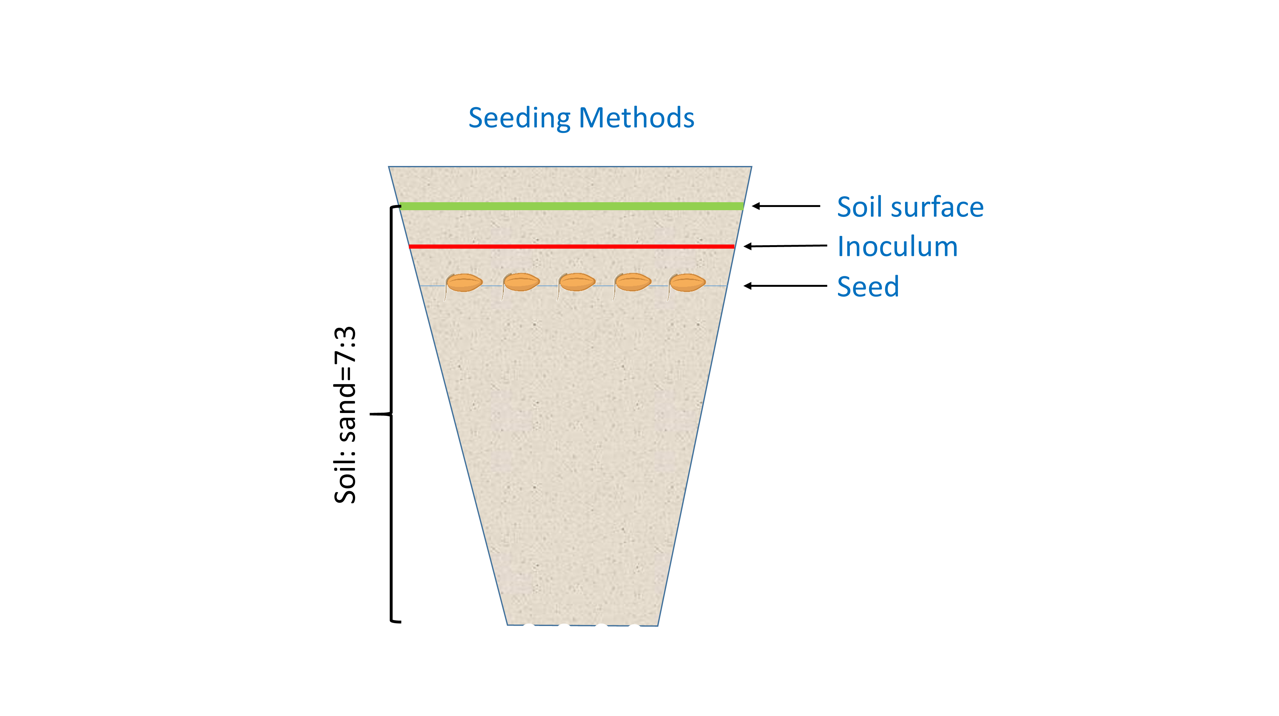

**Fig.S1** Schematic representation of *Fp* inoculation method used in the glasshouse experiments.

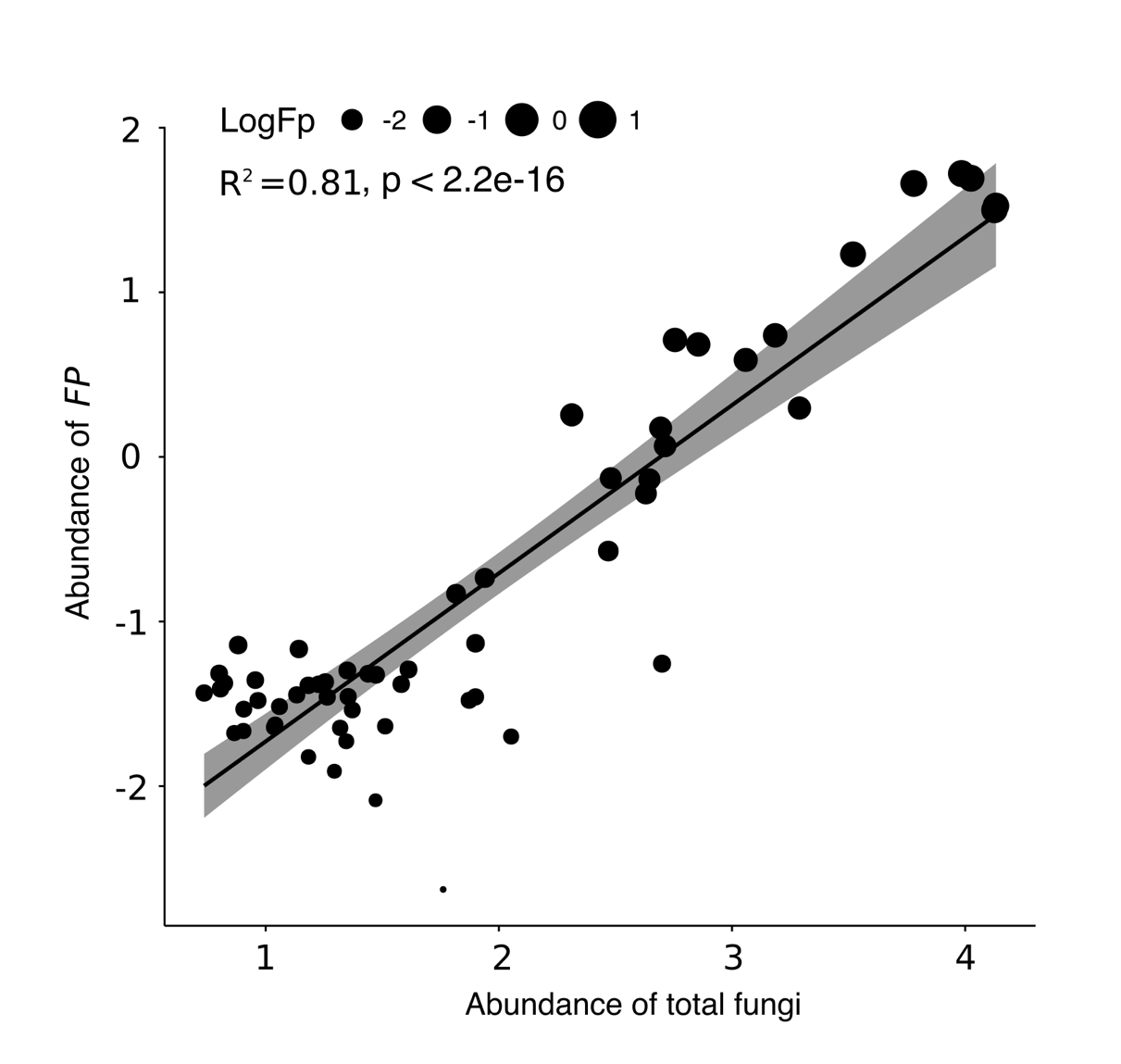

**Fig.S2** Correlation of *Fp* abundances with 18S rDNA copy numbers at the base of the wheat stem. Total fungi and *Fp* values have been log10 transformed (n=58).

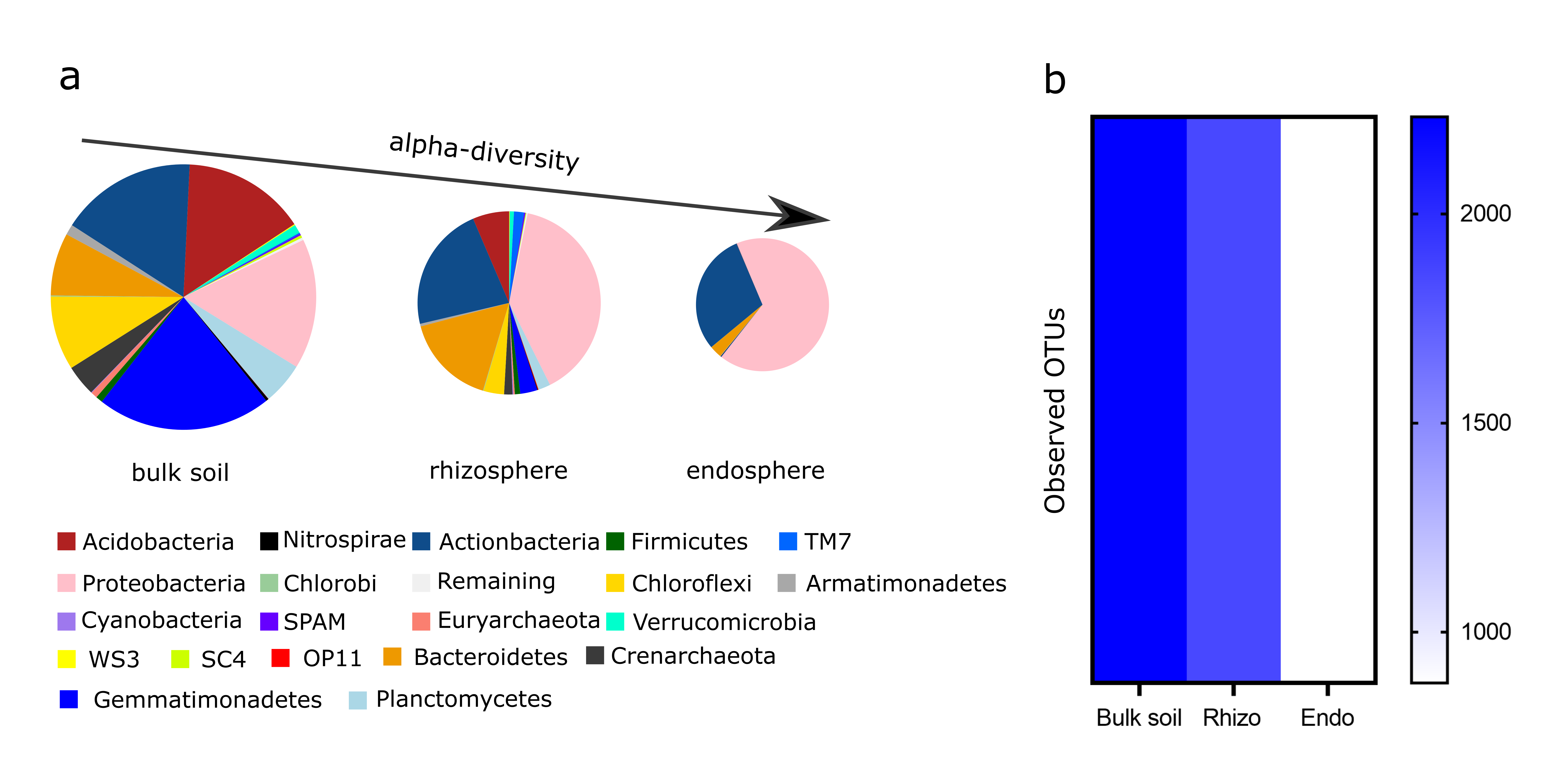

**Fig.S3** Gradient changes of the rhizosphere microbial communities. (**a**) Microbial composition changes from the bulk soil to the root endosphere, whereby the Proteobacteria increased in relative abundance while Acidobacteria and Gemmatimonadetes decreased. (**b**) A heatmap summarizing microbial diversity in the bulk soil, rhizosphere soil and the root endosphere.

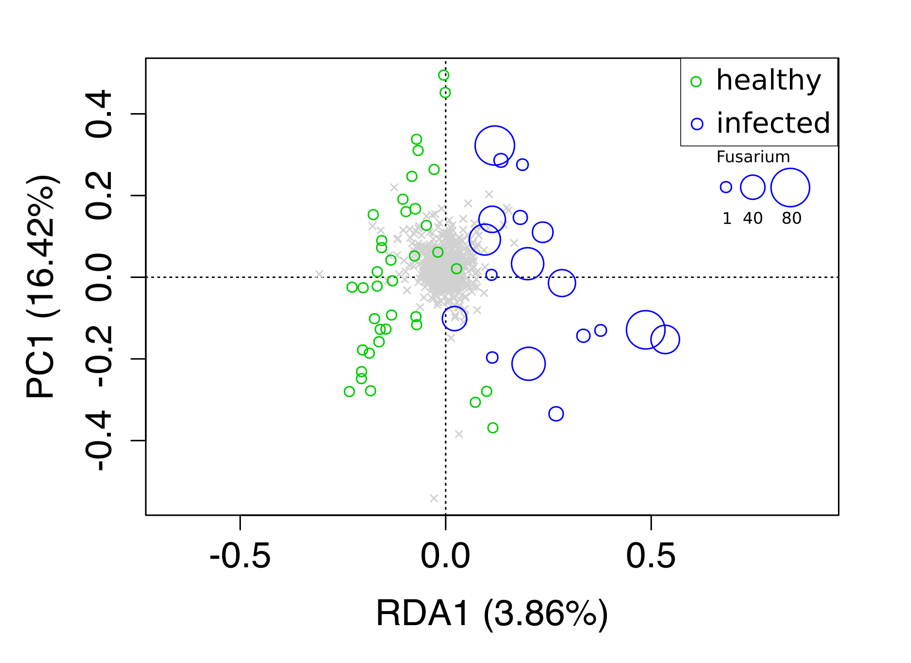

**Fig.S4** Redundancy analysis (RDA) summarizing the effect of *Fp* on fungal communities in the wheat rhizosphere soil. Circles in the panel are scaled to the abundance of *Fp* in wheat stems.

**a**

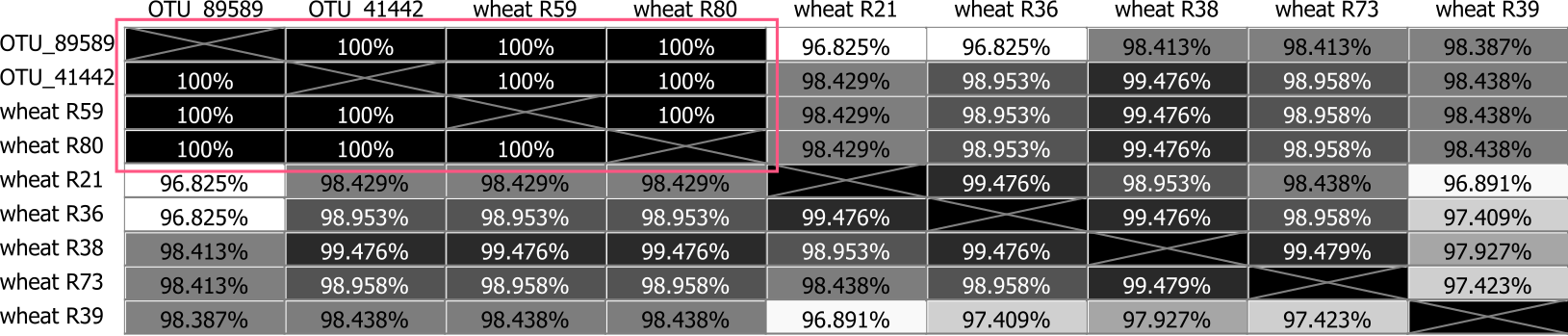

**b**

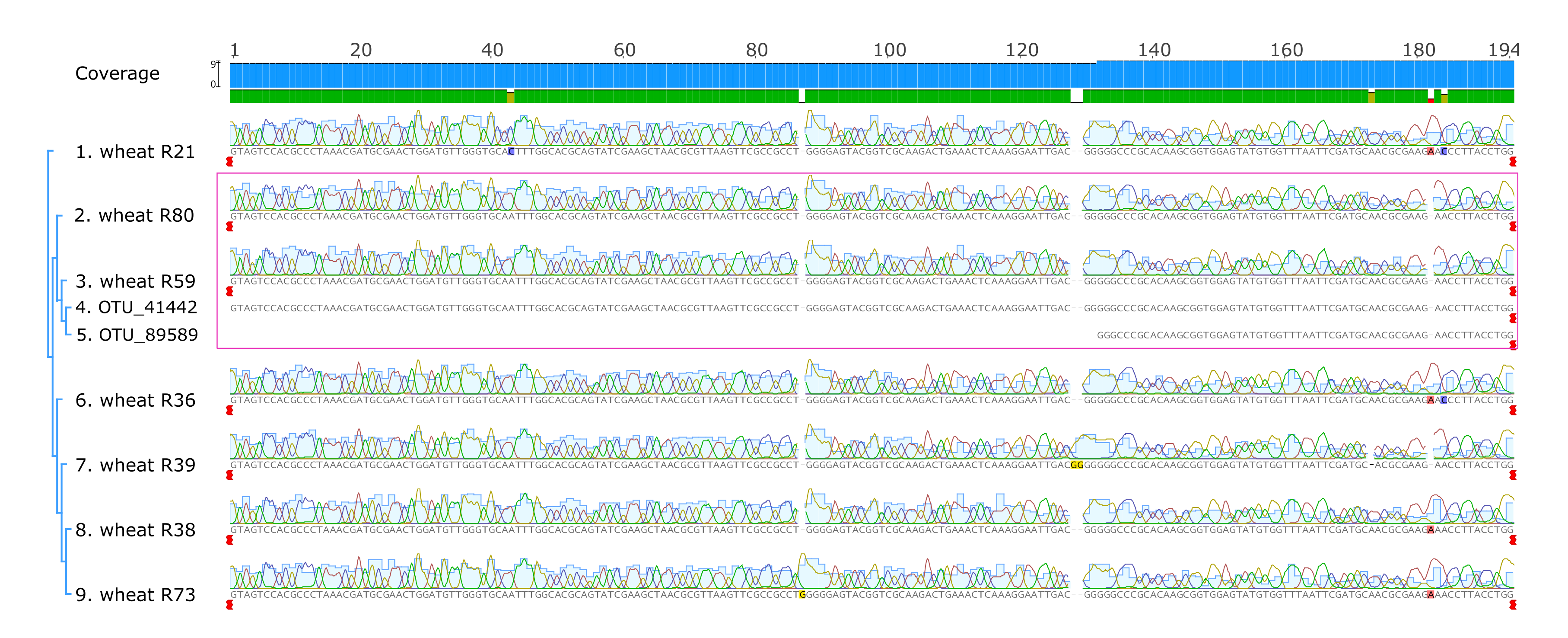

**c**

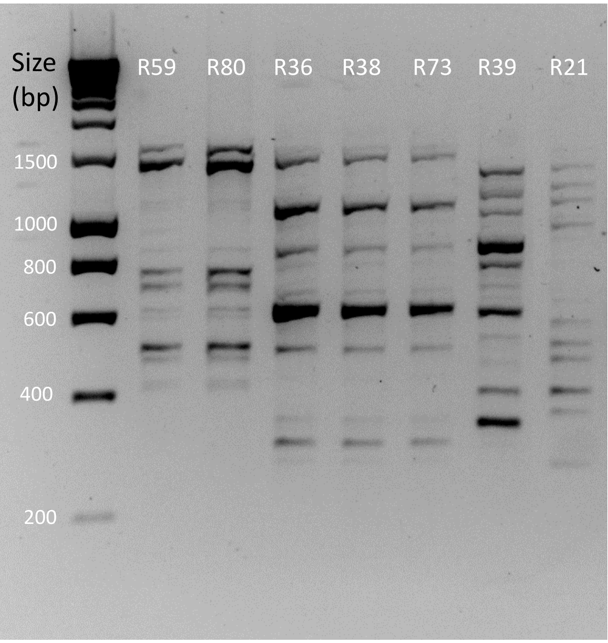

**Fig.S5** Phylogenetic analyses of seven *Stenotrophomonas* strains isolated from the wheat rhizosphere soil and roots. (**a**) Distance matrix of the 16S rRNA sequences of the isolated *Stenotrophomonas* strains and the OTUs 89589 and 41412. (**b**) Muscle alignment view of the 16S rRNA sequences of the seven *Stenotrophomonas* isolates and the 16S rRNA sequences of OTUs 89589 and 41412. Pink squares highlight the two strains (SR59 and SR80) that matched the sequences of the two OTUs. The analysis was performed using Geneious 10.1.3. (**c**) Fingerprint patterns of the seven bacterial strains typed using BOX-PCRs.

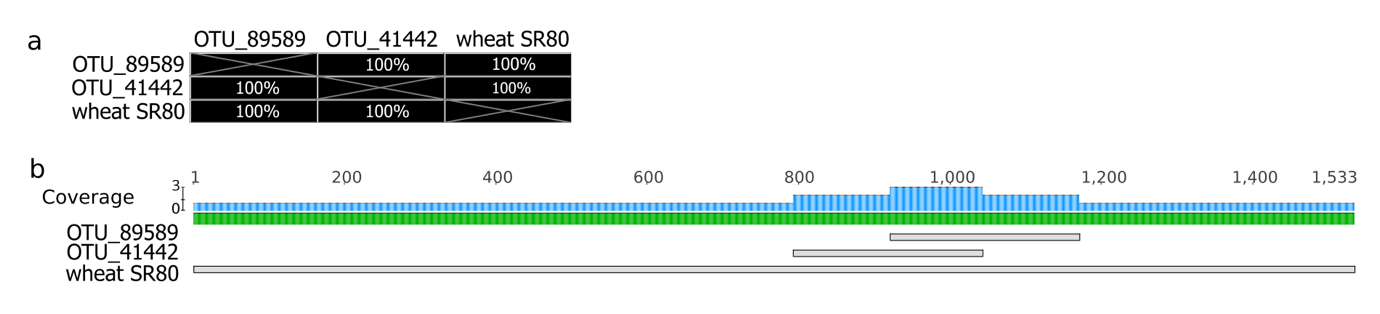

**Fig.S6** Alignment of the 16S rRNA sequences of the isolate SR80 and the two OTUs. (**a**) The distance matrix of the 16S rRNA sequences of SR80 (predicted by whole genome sequencing) and the OTUs 89589 and 41412. (**b**) Alignment view of the 16S rRNA sequences of SR80 with the OTUs 89589 and 41412. The analysis was performed in Geneious 10.1.3.

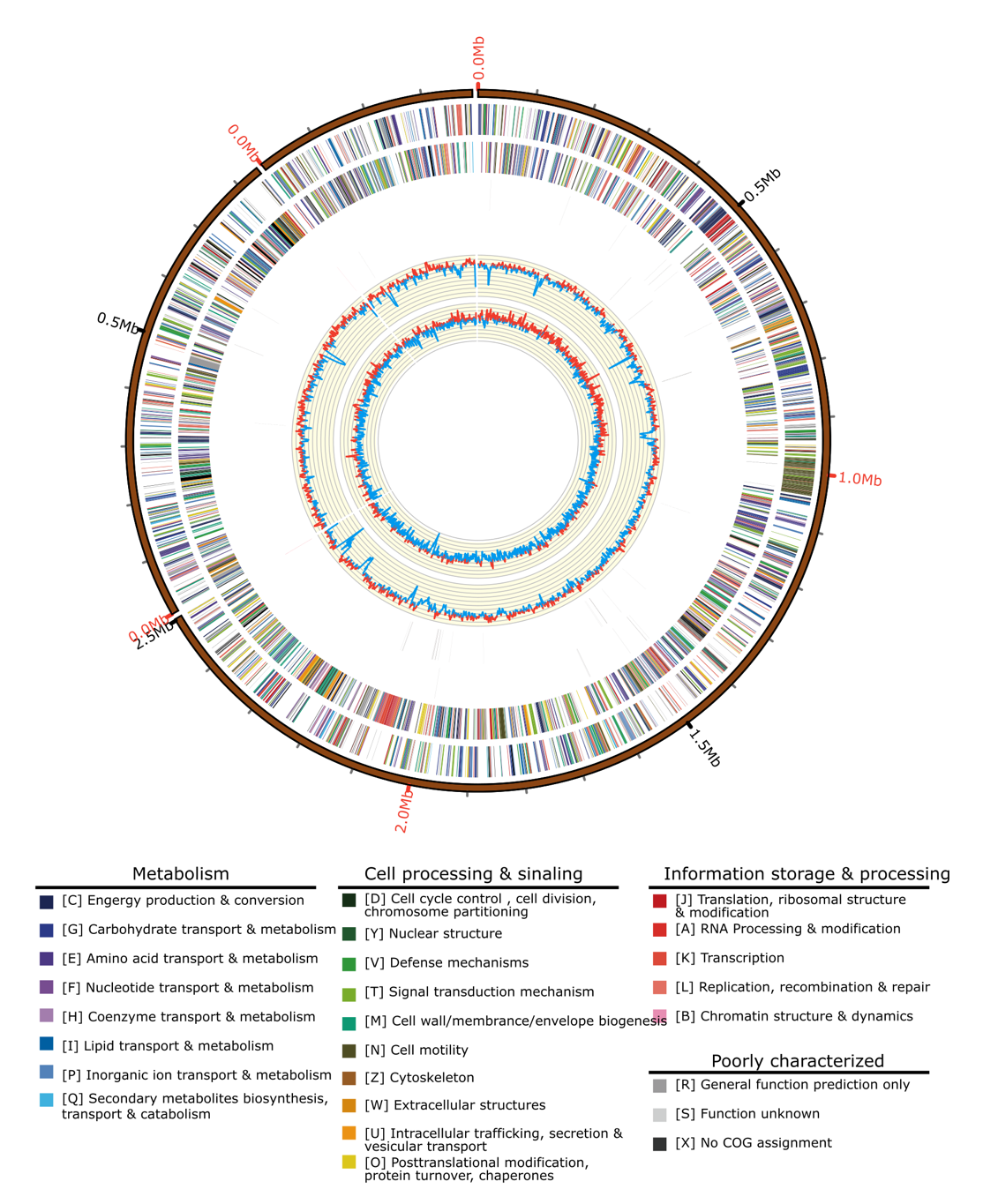

**Fig.S7** Circular map of the *Stenotrophomonas rhizophilia* R80 genome. The distribution of the circle from the outermost to the center is, (i) scale marks of the genome; (ii) protein-coding genes on the forward strand; (iii) protein-coding genes on the reverse strand; (iv) tRNA (black) and rRNA (red) on the forward strand; (v) tRNA (black) and rRNA (red) genes on the reverse strand; (vi) GC content; (vii) GC skew. Protein-coding genes are color coded according to their COG categories. The graph was generated using Circos v0.69 (<http://circos.ca/>).

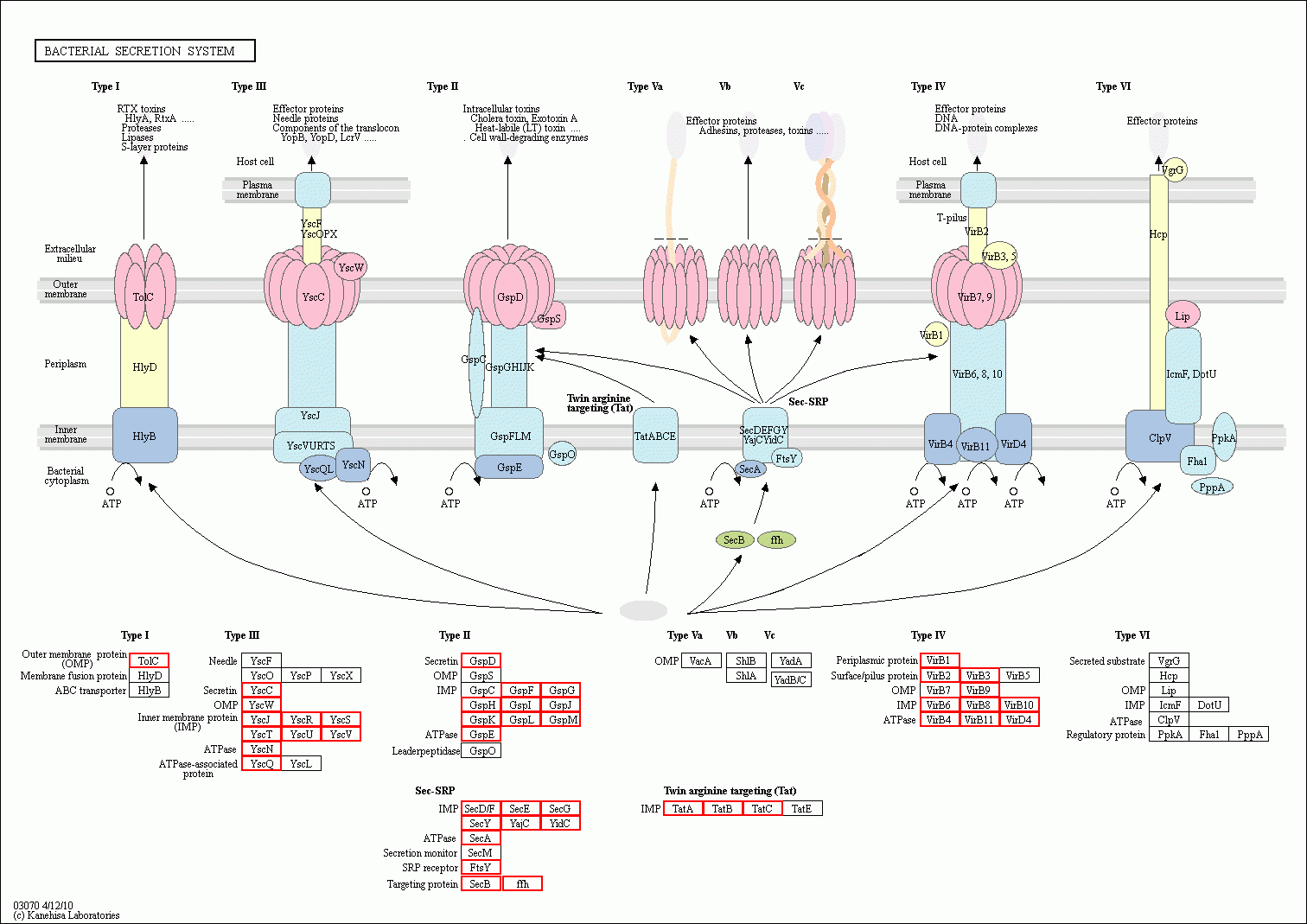

**Fig.S8** Prediction of secretion system encoding genes in SR80 genome. Red rectangles highlight those genes detected in the genome.

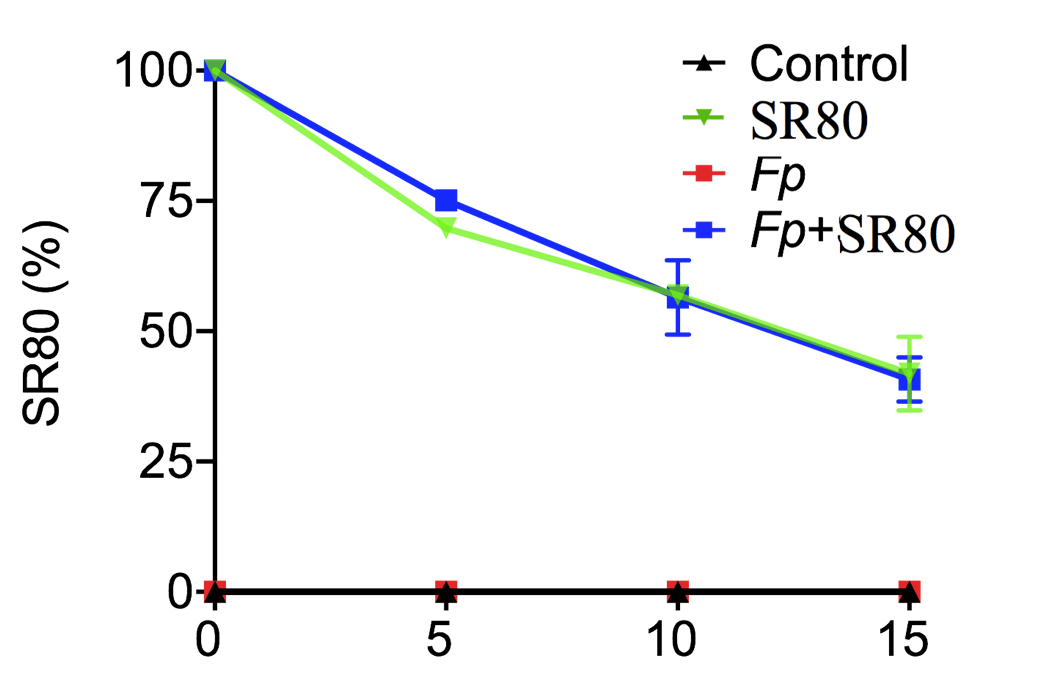

**Fig.S9** Changes of the relative abundance of SR80 in soils of the glasshouse experiment. Error bars represent standard errors of the mean (n=3).

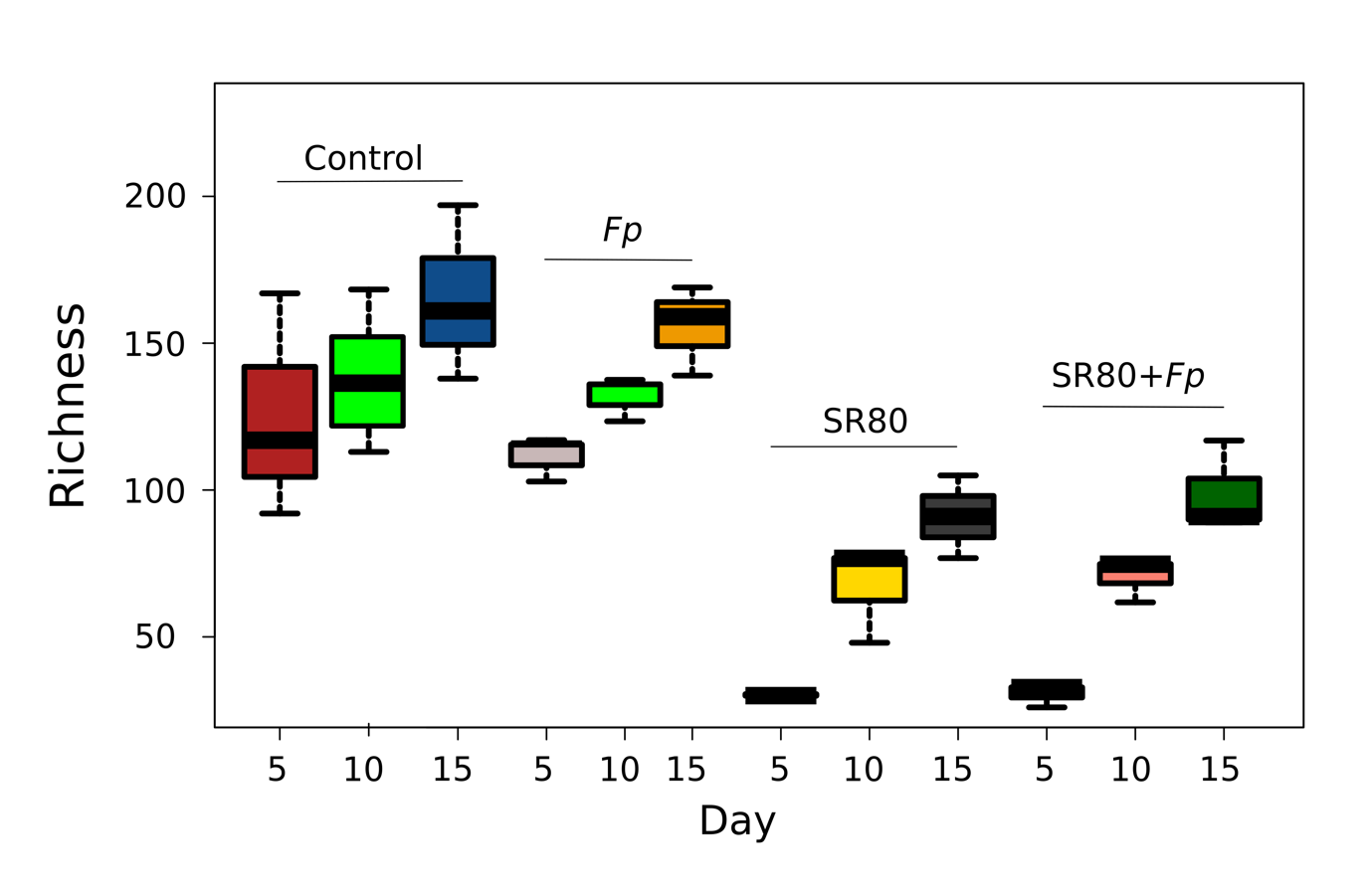

**Fig.S10** Changes of soil microbial diversity (observed species) under four treatments within 15 days. An increasing trend of the microbial diversity was observed for all four groups over time. Overall, the non-SR80 treated groups had higher diversity than the SR80 treated groups. Error bars represent standard errors of the mean (n=3).

**References**

**Andrews S 2010**. FastQC: a quality control tool for high throughput sequence data: Babraham Bioinformatics, Babraham Institute, Cambridge, United Kingdom.

**Bolyen E, Rideout JR, Dillon MR, Bokulich NA, Abnet CC, Al-Ghalith GA, Alexander H, Alm EJ, Arumugam M, Asnicar F. 2019.** Reproducible, interactive, scalable and extensible microbiome data science using QIIME 2. *Nature Biotechnol* **37**(8): 852-857.

**Callahan BJ, McMurdie PJ, Rosen MJ, Han AW, Johnson AJA, Holmes SP. 2016.** DADA2: high-resolution sample inference from Illumina amplicon data. *Nat Methods* **13**(7): 581.

**Cole JR, Wang Q, Cardenas E, Fish J, Chai B, Farris RJ, Kulam-Syed-Mohideen AS, McGarrell DM, Marsh T, Garrity GM, et al. 2008.** The ribosomal database project: improved alignments and new tools for rRNA analysis. *Nucleic Acids Res* **37**(suppl_1): D141-D145.

**Liu H, Carvalhais LC, Schenk PM, Dennis PG. 2017.** Effects of jasmonic acid signalling on the wheat microbiome differ between body sites. *Sci Rep* **7**(1): 41766.

**Liu H, Crawford M, Carvalhais LC, Dang YP, Dennis PG, Schenk PM. 2016.** Strategic tillage on a Grey Vertosol after fifteen years of no-till management had no short-term impact on soil properties and agronomic productivity. *Geoderma* **267**: 146-155.

**Liu H, Khan MY, Carvalhais LC, Delgado-Baquerizo M, Yan L, Crawford M, Dennis PG, Singh B, Schenk PM. 2019.** Soil amendments with ethylene precursor alleviate negative impacts of salinity on soil microbial properties and productivity. *Sci Rep* **9**(1): 6892.

**Rayment GE, Lyons DJ. 2011.** *Soil chemical methods: Australasia*: CSIRO publishing.
